# Human medial frontal neurons represent integrated semantic content and sensory evidence to inform decision-making

**DOI:** 10.64898/2026.11.06.764954

**Authors:** Tomas G. Aquino, Reidar Riveland, Robert Kim, Adam N. Mamelak, Chrystal M. Reed, Alexandre Pouget, Ueli Rutishauser, Nuttida Rungratsameetaweemana

## Abstract

Humans can reconfigure how they act on the world from a single verbal instruction. Central to this capacity is the ability to represent what an instruction specifies and to bring that content to bear on incoming sensory evidence. However, little is known about where in the human brain instruction content is encoded, how it combines with sensory evidence to produce a choice, and how this integration relates to behavior. Here, we characterized single-neuron and population responses recorded in the ventral temporal cortex, medial temporal lobe, medial frontal cortex and ventro-medial prefrontal cortex of neurosurgical patients performing a verbal instruction task. On each trial, a written instruction specified which stimulus feature was relevant and whether the correct response was to choose or to reject the stimulus that best matched it, so that the same image pair required opposite responses under different instructions. Only medial frontal populations jointly encoded the instructed task, and the target feature, whereas ventral temporal populations encoded stimulus features along axes that generalized across instruction conditions. Choice representation emerged first in ventromedial prefrontal and then in medial frontal cortex during stimulus presentation, before a motor response could be prepared. Shuffling instruction content selectively degraded prediction of medial frontal population activity, whereas shuffling stimulus identity selectively degraded ventral temporal activity. To ask how these representations could be computed, we trained a vision-language model with separable visual and linguistic streams converging on a recurrent decision module to perform the same task. Its visual embeddings predicted held-out ventral temporal activity and its recurrent activations revealed a task solution space that predicted medial frontal activity, and the strength of this alignment tracked behavioral accuracy. These results identify a cortical hierarchy in which medial frontal circuitry represents the content of verbal goals and acts on a ventral temporal substrate whose representational axes remain stable, and indicate that functional hierarchies discovered in artificial networks can serve as predictive models of human neural circuitry.

## Introduction

Goal-directed behavior is a crucial cognitive function, enabling individuals to produce adaptive decisions from goals. A uniquely human route to guide behavioral control is language: a verbal instruction can quickly specify new task rules, determine which sensory features are relevant, and determine which actions will be rewarded, often without extensive re-training on a new task set [1].

Previous foundational work has positioned frontal cortices as crucial centers for maintaining goals, exerting top-down control of perceptual decision-making, and flexibly integrating context-dependent evidence [2–6]. In parallel, the cortical hierarchy in higher visual areas, including inferotemporal cortex (IT) and ventrotemporal cortex (VTC), is known to parse out sensory information in increasingly complex bottom-up representations (e.g., objects and faces), which are utilized down-stream to enable efficient decision-making [7–11]. Recent results obtained in human single neurons have found visually selective cells in VTC for complex objects [12], building on previous results describing visual selectivity in human medial temporal lobe (MTL) [13]. We also determined that visual codes in VTC are reorganized depending on whether the current goals are known, modulated by medial frontal cortex (MFC) activity [14]. This suggests a rich interaction between MFC and VTC neural activity in integrating sensory and goal-related information during flexible decision-making in humans, building on work that demonstrated the role of human premotor and medial frontal cortices in encoding integrated decision variables [15], verbal instructions [16], and abstract goals [17], beyond motor outputs alone.

On the one hand, computational models have been proposed for the ventral visual stream (including VTC), via neural alignment with artificial networks [18, 19], and recurrent neural networks have been proposed as a model for context-dependent, multi-task decision making [5, 20, 21], mirroring the function of frontal cortices. On the other hand, the neural integration of instruction semantics with sensory evidence to produce a choice are not yet understood, especially at the rapid time scales at which behavioral control is reorganized by incoming instructions and stimuli.

To address this, we developed a verbal instruction decision task in which written instructions and visual stimuli jointly determine the correct response, while systematically manipulating task demands (feature selection across multiple stimulus domains), framing (pro versus anti instructions), and instruction availability during perception (early versus late instruction). Using intracranial single-neuron recordings across ventral temporal, medial temporal, medial frontal, and ventromedial prefrontal regions, we examined how sensory evidence and instruction semantics are represented and integrated during decision formation. In parallel, we trained a vision-language model (VLM) with separable visual and linguistic representations that converge in a recurrent decision module, enabling direct comparisons between artificial and biological representational hierarchies [21]. We show that ventral temporal populations primarily encode bottom-up stimulus features, whereas medial frontal populations emphasize instruction-dependent goal variables and choices, and that distinct model components predict neural variance in corresponding cortical stages. Together, these findings suggest a general computational principle for flexible behavior: hierarchical coordination that integrates goals that are communicated verbally with perception to guide decisions.

## Results

### Verbal instruction task

To examine how the human brain integrates instructions with sensory evidence to guide decisions, we designed a task in which written instructions and image pairs jointly determined the correct behavioral response (Fig. 1a). Each trial began with a 1-s fixation period, followed by the sequential presentation of two images (1 s each) drawn from four categories (faces, cars, animals, fruit). A written instruction specified the comparison rule, and a binary button-press response was required within 3 s. Instructions defined one of four task types: age comparison (faces/cars), color comparison (cars/animals), item-count comparison (animals/fruit), or identity comparison (faces/fruit). For age, color, and count tasks, the response indicated whether the first or second image satisfied the rule (e.g., “choose the oldest car”), whereas identity trials required a yes/no judgment about whether the two images depicted the same identity. Each session comprised 192 trials balanced across image categories and task types, organized into four blocks of 48 trials. Task framing varied between positive and negative instructions (e.g., “choose the oldest car” vs. “do not choose the oldest car”, see Extended Data Table 1). A key design feature of this task is disentangling bottom-up sensory perception from top-down goal-directed processing. Therefore, we presented the same set of stimulus images under different sets of instructions, or even in the absence of instructions. For this, instructions appeared either before image presentation (“early instruction”; Fig. 1a, left) or after the first image (“late instruction”; Fig. 1a, right), creating conditions in which some stimuli were viewed without prior task context.

**Fig. 1.**
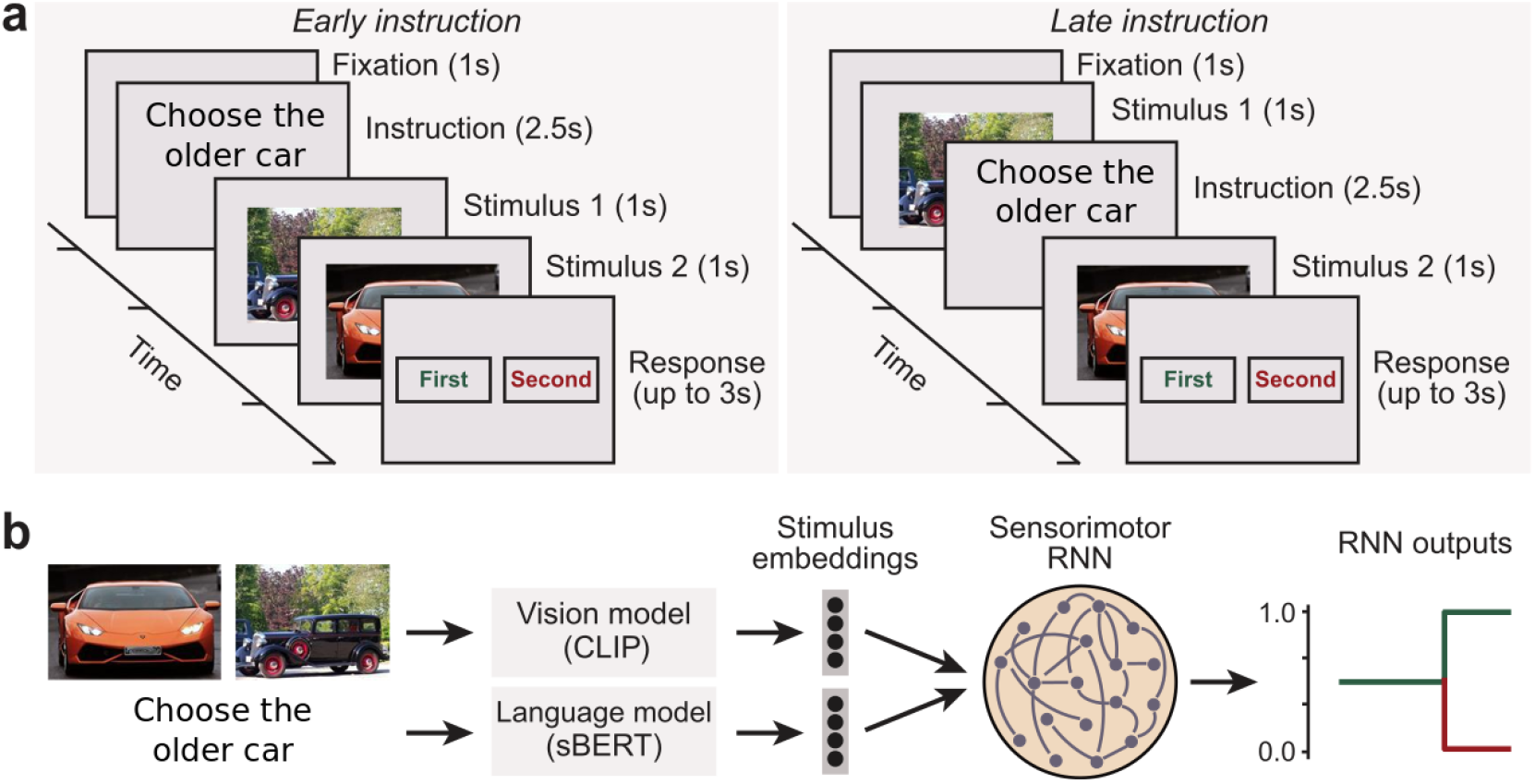
Task and model. **a**, Task. Patients saw an image pair and a natural language instruction in every trial, presented before the images in half the trials (early instruction, left), or between the images in the other half (late instruction, right). Subsequently, patients were prompted to respond according to the correct response to the instruction with a binary choice. **b**, Vision Language Model. The artificial agent consisted of parallel visual (CLIP) and linguistic (sBERT) streams, to generate embeddings for the images and instructions of each trial. Subsequently, these streams converged into a recurrent neural network (RNN) module, which produced a binary choice.

### A vision language model (VLM) and humans performed the verbal instruction task

To ask how verbal instructions and sensory evidence could be integrated to solve this task, we built a vision-language model (VLM) with separable visual, linguistic, and decision components (Fig. 1b). The VLM included fine-tuned vision and language modules, which generated embeddings that projected to a recurrent neural network (RNN) to generate binary decisions.

While the model’s performance was higher for trials containing face stimuli than other categories (Fig. 2a, *P <* 0.001 vs. all, two-sided Wilcoxon signed-rank test), it performed above 80% for all categories. Similarly, count and age tasks elicited better model performance, with age being higher than all the others (Fig. 2b, *P <* 0.001 vs. all, two-sided Wilcoxon signed-rank test). Additionally, pro-task framing trials elicited better performance than anti-task (Fig. 2c, excluding match trials since they disregard task framing, *P <* 0.001, two-sided Wilcoxon signed-rank test), while instruction order had no effect on performance (Fig. 2d, *P >* 0.05, two-sided Wilcoxon signed-rank test). To visualize the internal representations contained in separate modules of the VLM, we performed a principal component analysis (PCA, 3 dimensions) of the activations contained in the language module and observed how semantically similar instructions are closer in principal component space (Fig. 2e, e.g., “avoid the less colorful animal” and “select the creature with more colors”). To distinguish coding of semantic instruction meaning from coding of the visual appearance of letters on the screen, we utilized a pretrained vision model (see *Methods*) to create embedded representations of screenshots from verbal instructions exactly as displayed on the screen to patients. Qualitatively, instructions which had similar words and visual characteristics tended to cluster closer together, despite possibly having opposing meaning (Fig. 2f, e.g., “select the creature with fewer colors” and “select the creature with more colors”). We display a similar summary of image module embeddings in principal component space (Fig. 2g), as well as sensorimotor embeddings obtained from displaying the integrated RNN activations in principal component space (Fig. 2h). Qualitatively, the sensorimotor embeddings are sensitive to task goals, as different representation clusters arise depending on the magnitude of the target feature (low or high color/count/age), as well as on which stimulus is the correct answer to the trial’s instruction (first vs. second stimulus). It is possible to correctly decode several task factors (count, color, age, task type, target feature, *P <* 0.001 for all variables except count, which had *P <* 0.01, permutation test) from the RNN activations with a linear discriminant analysis (LDA, Fig. 2i).

**Fig. 2.**
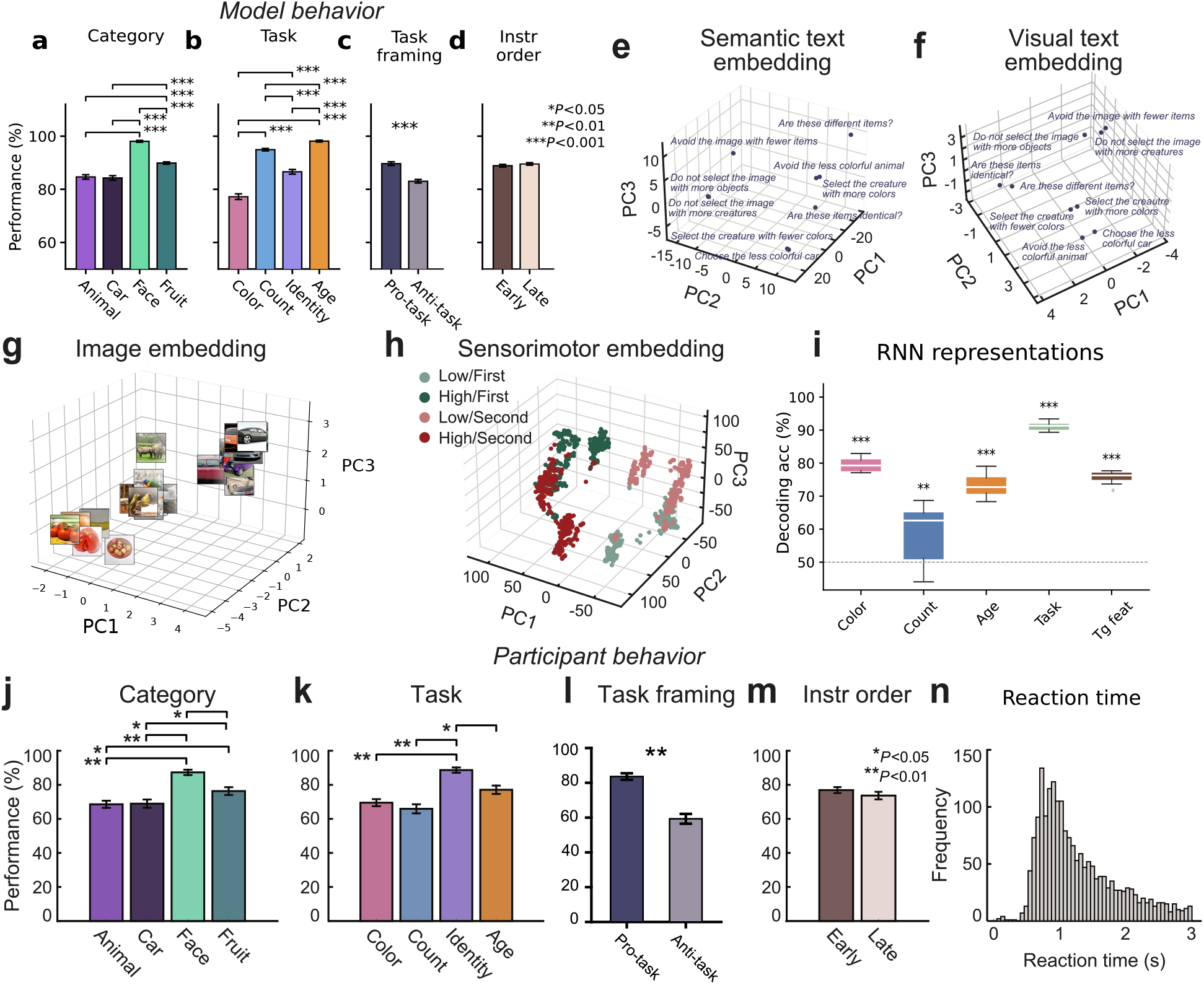
Vision Language Model and human participant performance. **a–d**, Model behavior. From left to right: dividing by stimulus category, task type, task framing, and instruction order. Error bars indicate s.e.m. (***: *P <* 0.001, **: *P <* 0.01, *: *P <* 0.05). **e**, Examples of text embeddings produced by the model, based on the semantic content of instructions. **f**, Same, for visual text embeddings based on the instruction screenshots seen by patients. **g**, Three principal components of CLIP ViT-B/32 image embeddings. Thumbnails are CC0-licensed stand-in images, embedded with the identical pipeline; the task stimuli themselves are not shown for copyright reasons. **h**, Visualizing task-based instruction structure in the model. **i**, Decoding stimuli and task representations from RNN module (***: *P <* 0.001, **: *P <* 0.01, permutation test). **j–m,** Participant behavior. From left to right: dividing by stimulus category, task type, task framing, and instruction order. **n,** Reaction time histogram across all sessions.

Patient performance on the task was above chance in all individual sessions (*P <* 0.001 for all patients, binomial test vs. 50% chance level). Trials containing face stimuli were performed at higher accuracy by patients (Fig. 2j, *P <* 0.01 vs. animals and cars, *P <* 0.05 vs. fruit, two-sided Wilcoxon signed-rank test), as well as identity matching trials (Fig. 2k, *P <* 0.01 vs. color and count, *P <* 0.05 vs. age, two-sided Wilcoxon signed-rank test). Similarly to the VLM, pro-task framing elicited higher performance than anti-task (Fig. 2l, *P <* 0.01, excluding match trials since they disregard task framing, two-sided Wilcoxon signed-rank test), while instruction order did not have an effect on performance (Fig. 2m, *P >* 0.05, two-sided Wilcoxon signed-rank test). Patients were able to perform the task within the stipulated time limit (3s) in the majority of trials (Fig. 2n).

### VTC and MFC neurons define a sensory/goal hierarchy

We recorded single neuron data from patients during epilepsy monitoring (see *Methods* for details on recordings and pre-processing). Following spike sorting, we obtained a total of 834 neurons across 11 patients: 137 in hippocampus (HPC), 143 in amygdala (AMY), 74 in dorsal anterior cingulate (dACC), 185 in pre-supplementary motor area (preSMA), 131 in ventromedial prefrontal (vmPFC), and 164 in ventral temporal cortex (VTC) (Fig. 3a).

**Fig. 3.**
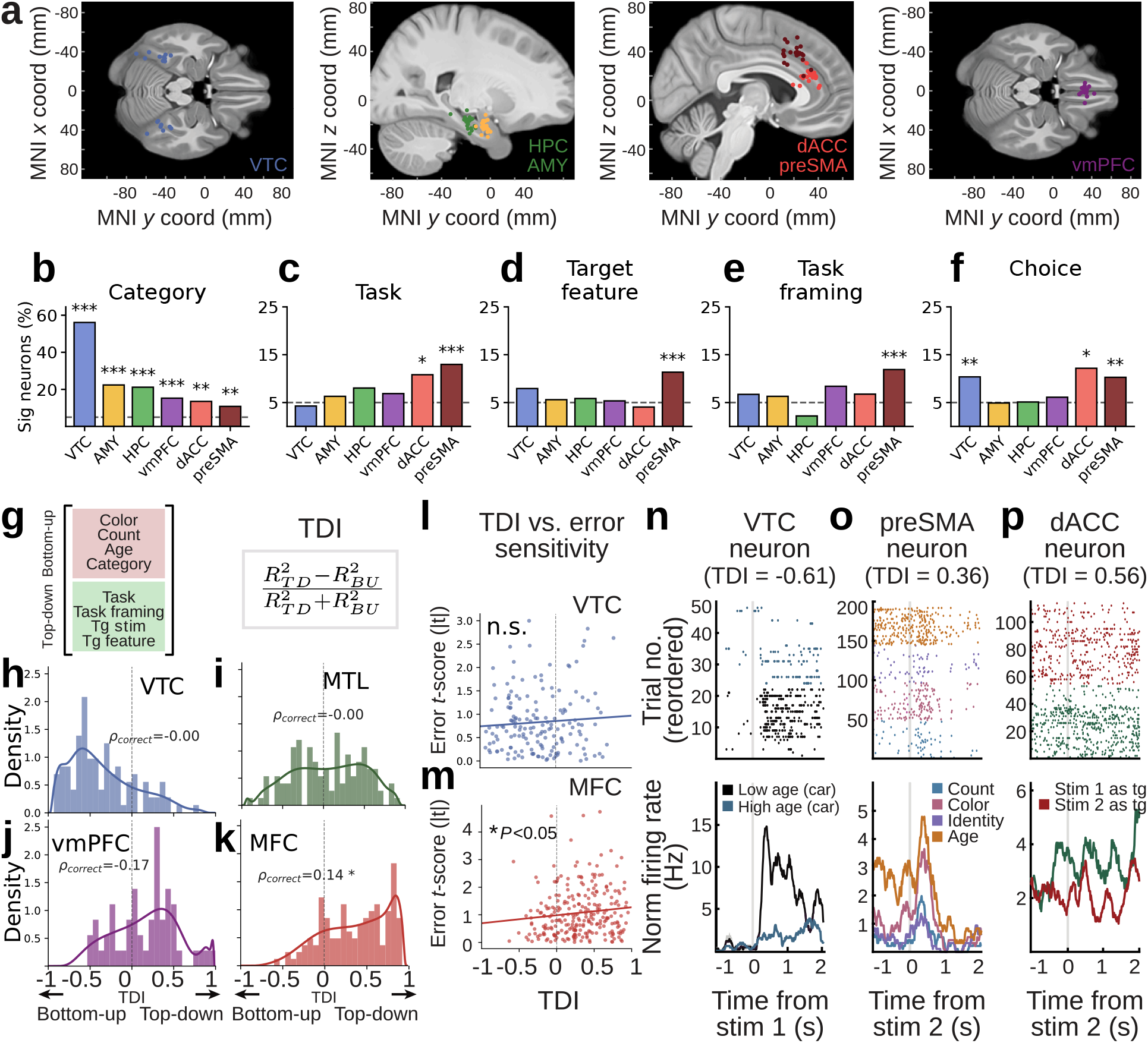
Electrodes, single neuron encoding, and top-down index. **a**, Electrode positioning in VTC, preSMA, dACC, vmPFC, hippocampus and amygdala. **b–f**, Proportion of neurons correlated during stimulus presentation with stimulus category, task type (color/age/count/match), target feature, task framing, target stimulus (***: *P <* 0.001, **: *P <* 0.01, *: *P <* 0.05, binomial test). **g**, Variables in the bottom-up and top-down blocks of the top-down index (TDI). **h–k**, TDI histograms for neurons with significant tuning to at least one of the tested variables, in VTC (h), MTL (i), vmPFC (j) and MFC (k). Inset: Pearson correlation between TDI and absolute *t*-score for correct versus incorrect coding. **l–m**, TDI versus error sensitivity in VTC (l) and MFC (m). **n–p**, Single neuron examples. From left to right: VTC stimulus feature neuron, preSMA task type neuron, dACC target stimulus neuron. Top: example neuron rasters. Bottom: smoothed peristimulus time histogram (PSTH). Shaded areas indicate s.e.m.

We first analyzed the tuning of each neuron individually in an encoding analysis, by correlating spike counts during stimulus presentation (pooling first and second stimuli) with task factors, correcting for reaction in the response phase as a proxy for trial difficulty (see *Methods* for encoding analysis). A significant number of neurons in all brain regions correlated with the visual category of a stimulus (face, car, fruit, or animal; Fig. 3b). Neuron count significance is determined via binomial tests (see *Methods*) compared to a chance level of 0.05.

The category variable had strong correlation with neural activity across brain areas (*P <* 0.001 for all brain areas, except dACC and preSMA, which had *P <* 0.001; VTC: 56.0%, AMY: 22.3%, HPC: 21.1%, vmPFC: 15.2%, dACC: 13.5%, preSMA: 10.8%). The task variable (comparing color, count, age, or identity), as determined by instructions, correlated with a significant number of neurons in dACC (10.8%, *P <* 0.001) and preSMA (12.9%, *P <* 0.001), Fig. 3c. The target feature (whether the current stimulus had a high or low level of the instructed feature) correlated with a significant number of neurons in preSMA (11.3%, *P <* 0.001), Fig. 3d. We report examples of MFC target feature coding neurons whose firing rate is a function of target feature axes for specific instructions (Extended Data Fig. 1a–c), along with an RNN node that also activates specifically depending on which target feature is displayed (Extended Data Fig. 1d).

Task framing (pro- or anti-task instructions, excluding match trials since they did not distinguish framing) correlated with a significant neuron count in preSMA (11.8%, *P <* 0.001), while choice correlated with firing rates in a significant amount of neurons in VTC (10.3%, *P <* 0.01), preSMA (10.2%, *P <* 0.01), and dACC (12.1%, *P <* 0.05), Fig. 3f. We also found that the activity of VTC neurons correlated with the age, count, and color features of stimuli (Extended Data Fig. 2a–c), whereas current task state (whether the current stimulus being visualized was the first or second), and instruction order (whether the current trial was an early or late instruction trial), had widespread correlation across brain areas (Extended Data Fig. 2d–e).

Put together, these results indicate a predominance of VTC in coding bottom-up, sensory related factors, as expressed in coding the visual categories of stimuli, whereas preSMA and dACC neurons more predominantly encode top-down factors related to instructions, task goals, and choice. To summarize the area-specific roles in encoding bottom-up or top-down variables, we created a top-down index (TDI, see *Methods*), which describes the relative correlation between neural tuning and top-down/bottom-up variables. We obtained TDIs for each individual neuron by using partial variance explainability in spike counts from top-down and bottom-up variables in a joint linear model and normalizing these factors to a [−1, 1] scale (Fig. 3g–k). Utilizing TDI distributions, VTC significantly displays more bottom-up coding than other regions (*P <* 0.001 vs. all regions, Kolmogorov-Smirnov test), whereas MFC displays significantly more top-down coding (*P <* 0.001 vs. all regions, Kolmogorov-Smirnov test). Additionally, we tested whether TDIs correlated with behavioral performance on the task and determined that top-down coding in MFC (but not in VTC) correlates with neural sensitivity to behavioral errors (Fig. 3l–m, MFC: *P <* 0.05, Pearson correlation test). These results further demonstrate the presence of a behaviorally relevant cortical hierarchy from VTC to MFC in encoding bottom-up versus top-down information during decision-making. For visualization purposes we included three example raster plots of neurons in VTC (Fig. 3n), preSMA (Fig. 3o), and dACC (Fig. 3p), respectively tuned to stimulus features, task instructions, and choice.

### Frontal neural populations represent task goals and choices, while VTC populations code visual stimulus features

To determine the populational dynamics of information representation during the task, we constructed pseudopopulations of neural activity across patients and performed SVM decoding with a linear kernel (see *Methods*), for several factors relating to stimuli and task goals. To simplify our decoding analyses, we pooled AMY and HPC together into a region called medial temporal lobe (MTL), as well as dACC and preSMA into a region called medial frontal cortex (MFC). Briefly, we assembled matrices containing spike counts across patients, from stimulus presentation windows, aligning trials across sessions that were matched in specific variables of interest (see *Methods*).

With this, we first decoded stimulus categories from the first stimulus presentation window (to avoid possible confounds with category information presented via instructions), obtaining prominent category decoding in VTC and MTL (Fig. 4a, *P <* 0.01, permutation test). Pooling first and second stimulus presentation windows, we also decoded whether the current stimulus was high or low in each visual feature, relative to its pair in each trial, resulting in binary age, count, and color features to be decoded. We successfully decoded all stimulus features from VTC (Fig. 4b–d, *P <* 0.01 for all, permutation test), as well as age (*P <* 0.01, permutation test) and color (*P <* 0.01, permutation test) from MTL. This further reinforces the prominent role of VTC in encoding sensory information pertaining to visual stimulus features.

**Fig. 4.**
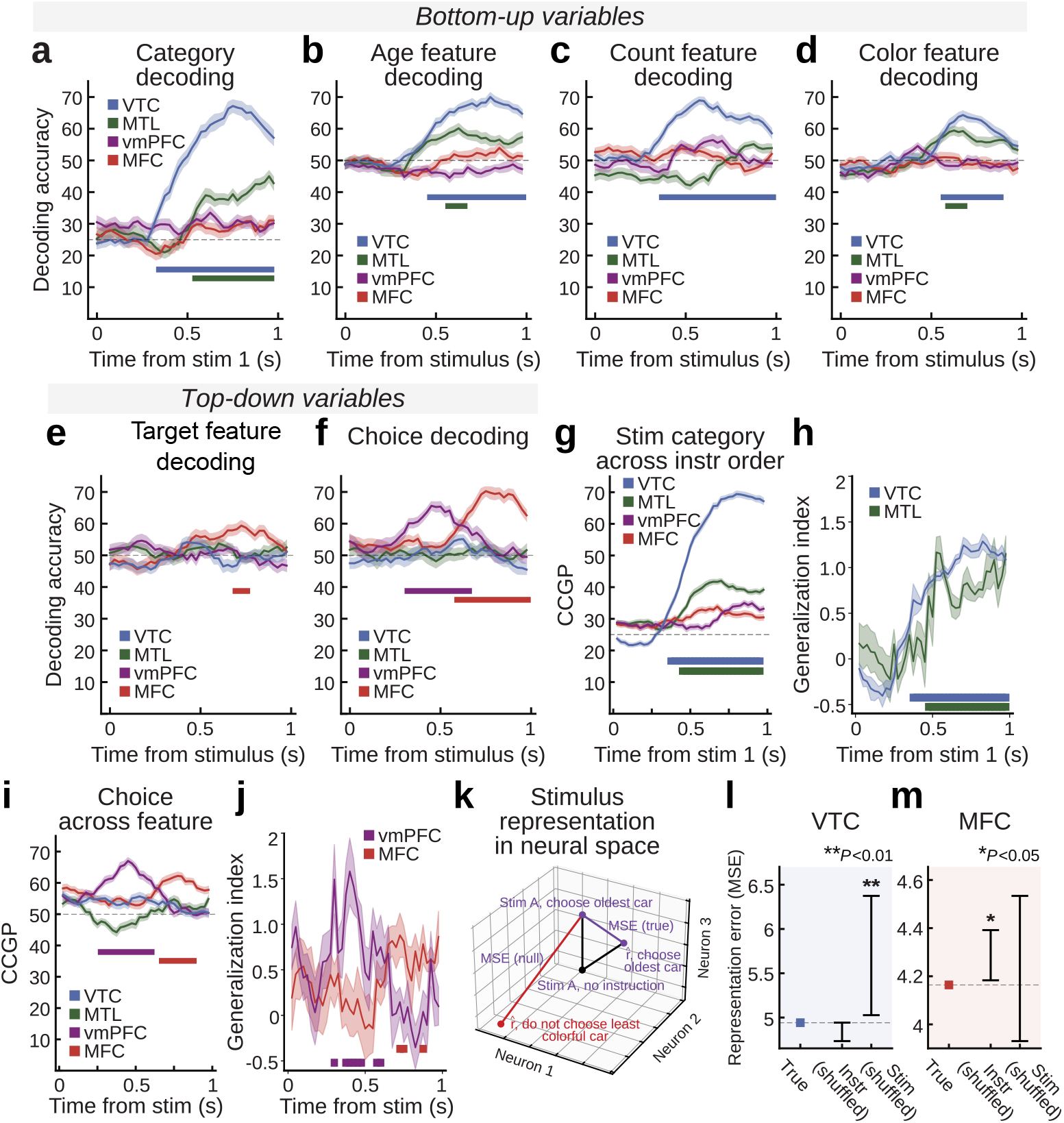
Populational representation of stimuli, task components, and choice. In panels a–j, shaded areas indicate s.e.m. and bars indicate significant time bins. **a–d**, SVM decoding during stimulus presentation. From left to right: stimulus category, age, count, and color. **e–f**, Decoding task related factors (left: target feature, right: target stimulus (choice)). **g**, CCGP for stimulus category across instruction order (early vs. late). Decoders were trained in early instruction trials and tested in late instruction trials. **h**, Generalization index for category across instruction order. Same train and test direction as g. **i**, CCGP for target stimulus (first vs. second) across target feature magnitude (low vs. high). Decoders were trained in low feature level trials and tested in high feature level trials. **j**, Generalization index for target stimulus (choice) across target feature. Same train and test direction as i. **k**, Schematic for a function which predicts stimulus representations conditioned on verbal instructions. Prediction accuracy is evaluated via mean squared error (true MSE) and compared to estimated representations with shuffled instructions (null MSE). **l**, Error in predicting out-of-sample stimulus representations from stimuli and instructions in VTC. The true error (dashed line) is compared with null distributions obtained by holding stimuli constant and shuffling instructions (Instr shuffled), or by holding instructions constant and shuffling stimuli (Stim shuffled). Larger MSEs imply worse representation accuracy. True values smaller than a shuffled distribution imply presence of information necessary for representation prediction. **m**, Same, for MFC (dACC and preSMA).

Subsequently, we decoded a binary regressor indicating whether features along the instructed axis were high or low, for each stimulus presentation (Fig. 4e). For instance, if the current instruction was to “choose the oldest car”, that defined car age as the relevant feature, and the regressor would be 0 if the currently viewed car was the newer one in its pair, or 1 if it was the older car in its pair. This regressor is a direct means of measuring a conditional interaction between instructed top-down task goals and visual stimulus features. We could successfully decode this variable from MFC (*P <* 0.05, permutation test).

Subsequently, we also decoded the choice patients made in each trial, during stimulus presentation. The choice regressor was a binary vector representing whether the chosen stimulus was first or second in the sequence, regardless of which button press was physically made at subsequent stages of the trial. At the population level, choice could be decoded from vmPFC and, subsequently, from MFC (Fig. 4f, *P <* 0.01 for both, permutation test). These results indicate that frontal regions, in contrast with MTL and VTC, play a larger role in representing the top-down, goal-related factors of task representation. This includes parsing which features are currently relevant and representing choices during stimulus processing, before any motor commands actually have to be executed.

To determine whether the geometry of category coding was affected by presence or absence of instructions, we utilized a decoding generalization approach. Briefly, we train linear kernel SVM decoders with the previously described procedure but restraining training sets to early instruction trials, freezing decoders, and restricting testing sets to late instruction trials (and vice-versa, results displayed in Extended Data Fig. 3a–b). This yielded a cross-condition generalization performance score (CCGP), which indicated that category decoding accuracy in MTL and VTC remained qualitatively stable across conditions (Fig. 4g), in comparison to within-condition decoding (Fig. 4a). To quantify how CCGP compares to within-condition decoding, we utilized a generalization index [17] (GI, Fig. 4h, see *Methods*), and determined that both in MTL and VTC category coding occurs in an abstract manner with respect to the presence of instructions, with GI values approaching 1 and significant GI in similar periods to within-condition decoding (permutation test, cluster corrected, *P <* 0.01). Similarly, we tested whether choice coding in MFC and vmPFC occurred in an abstract form relative to the targeted feature, training in low level of target feature and testing in high level of the target feature (and vice-versa, results shown in Extended Data Fig. 3c–d). That is, was the geometry of choice coding disturbed by which of the two possible features was the target of each trial? We observe successful cross-condition choice decoding in vmPFC and MFC (Fig. 4i, *P <* 0.01, permutation test), and periods with higher than chance choice generalization for MFC and vmPFC (Fig. 4j, permutation test, cluster corrected, *P <* 0.05). This suggests an abstract coding of choice with respect to target feature in vmPFC and MFC.

To determine whether instruction meaning representation was stable independent of wording, we performed a separate generalization analysis, in which we trained and tested a target feature decoder (linear SVM, same method as other decoding analyses), utilizing antonyms and double negatives to anchor meaning. Specifically, aggregating data across the entire stimulus presentation window (stim1, when instruction was available, and stim2), we trained the target feature decoder in trials of the first prompt variant (e.g. “Choose the most colorful car”) and tested in trials of the second prompt variant, which contained antonyms of the first prompt variant (e.g. “Choose the least colorful car”). Since task framing could be positive or negative (i.e. “do” versus “do not”) in both train and test sets, the same intended meaning of the instructions could appear in both sets, despite the first and second prompt variants containing antonyms. As such, we trained a decoder to detect the target feature in the first set, testing in the second set, and vice-versa. Bootstrapping a null distribution with 500 permutations (Bonferroni corrected for 4 brain areas) and averaging across the bidirectional generalizations, target feature generalization accuracy was 55.2% in VTC (*P <* 0.05, permutation test), 54.1% in MTL (n.s., permutation test), 56.5% in vmPFC (*P <* 0.01, permutation test), and 66.8% in MFC (*P <* 0.01, permutation test). These results confirm that coding of instruction meaning is most prominent in MFC and that its representation is persistent across different specific forms of wording through which the meaning is conveyed.

To further quantify the impact of verbal instructions on single-trial populational representations, we developed a method to predict single-trial stimuli representations given an [instruction, stimulus] pair (see *Methods*). Briefly, we trained a multi-layer perceptron to predict neural activity from stimuli while allowing an instruction embedding to gate this prediction. With this, we can predict the representation of a stimulus following an instruction relative to its representation in late instruction trials (Fig. 4k). To determine whether this prediction is valid, we compare the error of our estimated representation to the error of bootstrapped null representations, obtained by shuffling either the stimulus set or the instruction set. We subsequently compared the error in estimating representations versus instruction-null sets and stimulus-null sets to claim whether either of these factors was relevant in predicting neural representations. In VTC neurons, shuffling stimulus information disrupted representation prediction (*P <* 0.01, permutation test), while shuffling instruction contents had no effect (Fig. 4l). The opposite was true in MFC: shuffling instruction contents disrupted representation prediction (*P <* 0.05, permutation test), while shuffling stimuli had no effect (Fig. 4m). In conjunction with our single neuron encoding results, these analyses suggest a central role for MFC in encoding task goals and linguistic instruction contents, while VTC has a role in representing visual stimulus features, as expected. In addition, by leveraging cross-condition generalization, we demonstrated that the neural code in which patients’ choices are represented in MFC is abstract from the target features themselves, despite both variables being encoded in MFC simultaneously (Fig. 4e,f,i,j).

### Sensorimotor VLM embeddings explain human neural activity

With the multimodal VLM we developed for this task, we can propose a computational explanation for the activation patterns in human neural populations. This analysis expands on previous work which predicted activation patterns in the visual ventral stream of mammals utilizing convolutional neural networks as an explanatory model [19], extending this approach to the flexible decision-making domain. Concretely, after a task design is defined, it is presented to both humans and bespoke models, producing both neural firing rates *y* and artificial model representations for the task *X* (Fig. 5a). The network is shown the same trial sequences as patients, so during first stimulus presentation on a late-instruction trial, the decision RNN has not yet received any instructions, exactly as the patient has not. More generally, any information that is unavailable at the time of a presentation is likewise unavailable to the network and the regression models.

**Fig. 5.**
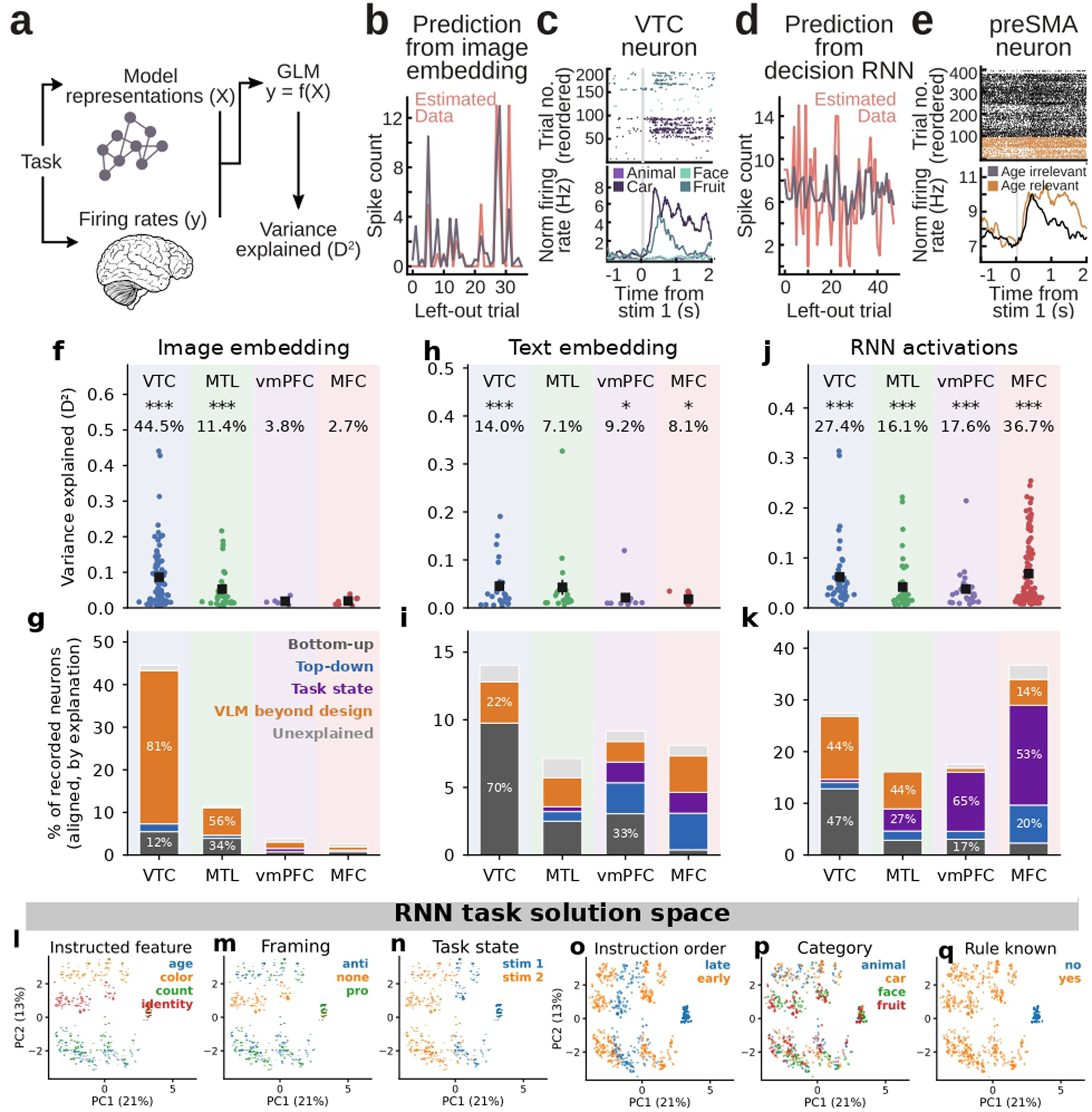
Vision language model discovers a task solution space that predicts human single neuron activity. **a**, Schematic for predicting variance in single neurons from activations in artificial models. VLM activations become latent space features for a Poisson-Gamma generalized linear model (GLM), which predicts spike counts in individual neurons. **b**, Example VTC category neuron whose activity is predicted out-of-sample by the VLM. Real and estimated spike counts in left out trials. **c**, Raster and PSTH for this neuron. **d–e**, Same, for a task-related preSMA neuron correlating with whether age was relevant or not. **f**, Variance explained (*D*^2^) in neurons whose activity was significantly predicted out-of-sample by CLIP image embeddings, after a permutation test. Box and whiskers indicate mean and s.e.m. **g**, Post-hoc single neuron analysis for the significant neurons above, displaying the best explanation for each neuron’s variance, comparing CLIP image embeddings as an explanation versus top-down, bottom-up, and task state factors. **h–i**, Same, for predicting neural activity from sBERT text embeddings. **j–k**, Same, for predicting neural activity from RNN activations. **l–q**, Task solution space found by the RNN, represented as two top principal components from RNN activations, separating trials by instructed feature, task framing, current task state (first vs. second stimulus), instruction order (early vs. late), stimulus category, and whether the current trial instruction was known or not.

Our final goal is to predict neural activity patterns using artificial model activations as inputs. Before achieving this, these representations can be transformed in intermediate steps, such as reducing their dimensionality or aligning them into a shared latent space. In this case, we simply reduced the dimensionality of the model representations into a 20 dimension space via principal component analysis (see *Methods*). Subsequently, we fit Poisson-Gamma generalized linear models (GLM) to represent neural spikes as a function of artificial model activations stemming from CLIP image embeddings, sBERT text embeddings, and RNN activations. To quantify our ability to predict neural activity from model activations, we deployed an out-of-sample variance explained metric (*D*^2^ for the Poisson-Gamma models), obtained in left-out trials. We compared this metric for each neuron to its null counterpart, obtained by shuffling *y* and bootstrapping a null distribution for *D*^2^. We display two examples of neurons whose activity is predicted out-of-sample through this procedure, in VTC (Fig. 5b–c) and preSMA (Fig. 5d–e). While the VTC neuron was tuned to visual stimulus categories and was predicted by image embeddings from the VLM, the preSMA neuron displayed more complex activation patterns that were predicted by RNN activations, firing more in trials where age was the instructed feature, consistent with tuning to the instructed task.

To summarize how image embeddings explain variance across neural populations, we plotted the percentage of neurons in each brain area whose *D*^2^ metric is above 99% of their null counter-parts. We observe a significant number of neurons explained by image embeddings in VTC and MTL (*P <* 0.001, binomial test) (Fig. 5f). Post-hoc, we investigated this subset of neurons that was considered significant according to the variance explained metric. We tested whether each of these significant neurons was better explained by a linear combination of the known experimental design factors (Fig. 3, bottom-up factors, top-down factors, task state variables), or whether the VLM provided a better explanation to each of these neurons above and beyond known task design features. Concretely, this tests whether the VLM provides any additional insight once the linear task factors are in the explanatory model (Fig. 5g,i,k; full breakdown by variable in Extended Data Fig. 4a–c).

In VTC, most neurons aligned with CLIP image embeddings were better explained by the visual component of the VLM than by a model containing the known task design factors (Fig. 5g). This indicates that representations contained in CLIP image embeddings are parsing extra information that is relevant for predicting visual neural representation. This is consistent with the existence of a multidimensional visual code in human VTC that is extractable from representations of visual stimuli derived from deep neural networks [12]. In MTL, the largest share of image-aligned neurons was also better explained by the VLM (56%), while a considerable portion (34%) was explained directly by bottom-up factors, largely category tuning (Extended Data Fig. 4a). We also sought to determine whether the alignment between artificial and biological representations is predictive of behavioral performance on the task. For this, we developed a trial-wise deviance metric (see *Methods*) to compare the average performance of neural-artificial alignment in correct versus incorrect trials, as determined by patients’ choices during the decision phase. The alignment between neurons and CLIP image embeddings was not significantly predictive of behavioral performance in any of the tested brain areas (Extended Data Fig. 5a).

Similarly, we tested whether text embeddings from the VLM explained neural variance (Fig. 5h) and observed a significant count of explained neurons in VTC (14.0%*, P <* 0.001, binomial test), and at smaller proportions in vmPFC (9.2%*, P <* 0.05, binomial test), and MFC (8.1%*, P <* 0.05, binomial test). Post-hoc, the most parsimonious explanation for the majority of these VTC neurons is that they were tuned to stimulus category information, which is also conveyed via text (Extended Data Fig. 4b). We also determined that the alignment between text embeddings and MFC neurons was predictive of behavioral performance (Extended Data Fig. 5b, *P <* 0.05, one-sided *t*-test), indicating that medial frontal neural representations tend to be better predicted by VLM representations in trials which are behaviorally correct.

Finally, we performed the same analyses aligning RNN activations in the VLM and neural spikes (Fig. 5j), and determined that all brain regions had a significant number of neurons explained by the model (*P <* 0.001, binomial test). However, the region with the highest prevalence of neurons explained by the RNN activations was MFC (36.7%*, P <* 0.001, binomial test), out of which 53% were primarily neurons correlated with current task state (first vs. second stimulus) and instruction order (Fig. 5k). Still, 14% of MFC neurons whose activity correlated with RNN features were better explained by these features than by a model containing the known task design factors. In VTC, this proportion is 44% (Fig. 5k), which is similar the proportion of VTC neurons primarily explained by bottom-up factors, mostly category neurons (Extended Data Fig. 4c). RNN-neural alignment correlated with behavioral performance in vmPFC (*P <* 0.05, one-sided *t*-test) and MFC (*P <* 0.05, one-sided *t*-test), establishing the behavioral relevance of the VLM analysis for the representation of these brain areas (Extended Data Fig. 5c).

Put together, these results suggest that the VLM discovers a behaviorally-relevant task solution space which explains both sensory and decision-making activity in human single neurons, above and beyond what can be explained linearly by task design factors alone. To further characterize this task solution space, we visualize RNN activations along its two principal components, and determine that the RNN provides separable representations for different task types, task framing, task states, instruction order, stimulus category, and whether the current instruction was already known or not (Fig. 5l–q). For completeness, we also report visualizations derived from the CLIP and sBERT embeddings alone (Extended Data Figs. 6, 7), and display examples of neuron rasters which are explained by the VLM’s task solution space, plotted with respect to RNN principal components (Extended Data Fig. 8a–b).

Finally, in the task solution space, representations occupy an expanded volume once instructions are known, compared to before the instruction (Fig. 5l–q). We asked whether the same holds in the neural data by comparing the first stimulus of early-instruction trials with the first stimulus of late-instruction trials. These are the same images in the same serial position, differing only in whether an instruction preceded them. In MFC, the instruction-related spread of the representation (*P <* 0.01, permutation test) and its total occupied volume (*P <* 0.01, permutation test) were larger after the instruction than before it (Extended Data Fig. 9a–b, see *Methods*, *Expansion of the stimulus representation after the instruction*). The RNN showed the same pattern (Extended Data Fig. 9c–d). Before the instruction, presentations are labeled by an instruction that the patient has not yet seen, so the instruction-related spread is expected to be zero at that point. This analysis therefore shows that MFC populations carry the instruction once it has been given, in agreement with the model.

To control for the possibility that neurons in each brain area might be better explained by the visual aspect of instructions rather than semantic content, we performed a model comparison analysis for each neuron, comparing their cross-validated *D*^2^ between fits with the linguistic VLM embeddings versus fits with a transformer-based visual model of instruction panel screenshots (see *Methods*). Post-hoc, we selected the subset of neurons which were significantly explained by text embeddings for this model comparison analysis (Fig. 5i). In comparing the difference between semantic-based versus screenshot-based *D*^2^, VTC neurons collectively had higher *D*^2^ for the screenshot-based model (*P <* 0.05, two-sided, one-sample *t*-test, false discovery rate corrected across brain areas). Out of the 23 VTC neurons that were significantly explained by text embeddings (Fig. 5i), 18 (75%) were individually better explained by the screenshot model. Other brain areas did not have a significant difference between the screenshot versus semantic-based models. This control analysis suggests that text-based VTC neuron explainability relies more on visual features of presented letters and sentences than on the semantic and linguistic content of instructions themselves.

Put together, these findings indicate that the VLM is a viable method for predicting activation patterns in human neural populations during decision-making, based on its internal task solution space, and for capturing variation patterns tied to structural task factors such as instruction order and task state, providing insight for activity in neurons that were not previously understood based on linear combinations of task design factors alone.

## Discussion

We asked how the human brain maps a verbal instruction onto a perceptual decision, by recording single neurons across ventral temporal, medial temporal, medial frontal and ventromedial prefrontal cortex while patients perceived stimuli either in the presence or absence of instructions, and by training a vision-language model (VLM) to perform the same task. Three findings are, to our knowledge, new. First, medial frontal neurons integrate bottom-up stimulus features with the top-down goal set by the instruction: they encode a stimulus feature only when that feature is instructed, and they encode it along a target axis whose sign is set by the framing of the instruction, so that the same feature value of the same stimulus drives opposite responses under a pro- and an anti-instruction (Fig. 3d, Fig. 4e). Second, this integrated representation is predicted by a network that was trained to perform our multi-task battery, given that recent advances in NLP make it possible for the first time to model how humans follow linguistic instructions, utilizing artificial neural networks. We found that, by training on an identical task set, model activations are predictive of human single neuron activity in both sensory and decision brain areas. Out of the 834 recorded neurons, the VLM explained additional variance in 120 (14.3%, VTC: 63/164, MTL: 33/280, vmPFC: 5/131, MFC: 19/259), which could not previously be explained before from task design factors alone. The network’s decision layer defines a low-dimensional *task solution space* in which every stimulus-instruction pairing has a position (Fig. 5l–q), such that VTC and MFC neurons correlate with coordinates in that space. The degree of neural-RNN alignment in MFC and vmPFC neurons was also predictive of accuracy on the task, indicating the behavioral relevance of internal representations established in the VLM. Third, since every stimulus was seen both before and after an instruction, we could show how instructions re-map stimulus representations: the pre-instruction representation is compact and visually organized, and it expands along instruction-dependent directions once the rule is known, in MFC, as in the model (Extended Data Fig. 9), but not in VTC. We also reproduce a number of well-known results on cortical hierarchy between ventral temporal and medial frontal cortices during sensory processing and goal-directed decision making, in this new context of bottom-up/top-down integration.

Specifically, several of our observations reproduce established findings and consolidate them at single-neuron resolution in humans. Category and continuous-feature selectivity in VTC, and the fact that image-derived embeddings explain a large fraction of VTC neurons, replicate decades of work on macaque inferotemporal cortex [11, 22, 23] and its correspondence with human ventral temporal cortex [9, 12, 18, 19]. Category selectivity in the medial temporal lobe reproduces earlier human single-neuron reports [24, 25], and the encoding of the instructed feature, framing and upcoming choice in medial frontal neurons is consistent with rule and goal coding in primate prefrontal cortex [2, 3, 5, 26] and with human MFC recordings during conflict, memory-based choice and performance monitoring [15, 17, 27–29]. Most of the predictive power of the model in the frontal cortex is carried by task state and top-down variables, but we demonstrate there is an additional fraction of variance that can be better explained by the VLM’s task solution space.

Goal coding in the frontal cortex is well established, but a goal signal on its own does not say how a stimulus is evaluated under it. The central result of the single-neuron encoding analysis is that MFC neurons carry the evaluation itself. First, feature coding in MFC was relevance-gated: we discovered a population of MFC neurons whose activity correlated with the magnitude of the currently targeted feature, regardless of its sensory aspects (Fig. 3d, Fig. 4e). Conjunctive, mixed-selective coding of this kind has been described in primate prefrontal cortex and hippocampus as the substrate of context-dependent readout [5, 30, 31], and feature-based selection signals have been localized to frontal sources in the macaque [32]. Our results show that in humans this integration is carried by medial frontal neurons during stimulus processing, that it is specific to the instructed feature, and that it is signed by the verbal framing of the rule. None of these properties was present in VTC, where feature coding was neither gated by relevance nor signed by framing.

Category coding axes in VTC generalized between early- and late-instruction trials (Fig. 4g–h), which shows that these axes are preserved whether or not an instruction is present. This analysis does not test whether the content of the instruction changes VTC codes. The two analyses that do test content found no effect in VTC. Shuffling instruction content did not disrupt the prediction of VTC activity (Fig. 4l), and the VTC representation showed no significant instruction-related spread or expansion after the instruction (Extended Data Fig. 9). These are null results, and they do not rule out instruction-dependent changes in VTC. Indeed, the companion study found that visual codes in VTC are reorganized depending on whether the current goals are known [14]. Our results are therefore compatible with the literature reporting task- and attention-dependent modulation of inferotemporal responses in the macaque [33–35] and of human ventral temporal cortex measured with fMRI [36]. The stability of the category axes also agrees with previous work in which task context altered category representations in prefrontal but not in high-level visual cortex [37]. The medial temporal lobe behaved like VTC in the category generalization and expansion analyses, consistent with its visual alignment being carried by category. We urge caution in interpreting the absence of instruction effects in VTC, since it might be overturned with larger population recordings.

Additionally, both vmPFC and MFC represented choice at the population level, but only MFC showed the relevance-gated, target feature coding described above. MFC contained more neurons explained by the task solution space beyond the task variables, and only MFC showed the representation expansion seen in the VLM after the instruction. On the other hand, vmPFC was dominated by task-state coding (Fig. 5k) and its choice geometry was less disrupted when the relevant feature changed. This places vmPFC at a more abstract decision stage and MFC closer to rule implementation, where the choice remains bound to the currently relevant features. The ordering agrees with human single-neuron evidence that preSMA represents decision components while vmPFC codes value along the direction of choice [15] and with the dissociation of attention and decision computations across primate prefrontal regions [38].

Finally, our work builds on past NeuroAI literature utilizing artificial networks as explanatory models of the ventral visual stream [18, 19, 39] and of language comprehension [40, 41], as well as recurrent networks which have been proposed as models of flexible, instruction-dependent computation [20, 21, 42–45]. Our novel contribution is that our VLM supplied a task solution space in which stimulus, instruction, framing and the state of knowledge about the rule are arranged jointly, defined for every stimulus-instruction pairing. Using this space as a predictor uncovered neurons along the cortical hierarchy whose activity follows the integrated representation between stimuli and goals, the signature one expects of a conjunctive code for “this feature of this stimulus under this rule” [30, 31]. Consistent with this space, MFC representations of the same images acquired instruction-related structure once the instruction was known.

To conclude, verbal instructions determine contextual goals that interact with sensory representations to yield decisions. Our recordings locate that interaction: ventral temporal cortex provides a sensory representation whose category axes generalize across the presence and absence of an instruction, and medial frontal cortex re-maps the same stimuli into an instruction-dependent representation that a task-trained vision-language model predicts beyond what factorial description of the task can. The correspondence between the model’s solution space and medial frontal activity suggests that representations discovered in networks trained on a task can serve as quantitative hypotheses about the geometry of human neural codes.

## Acknowledgments

We are deeply grateful to the patients and their families for their willingness to participate in this research. We thank the staff of the epilepsy monitoring unit at Cedars-Sinai Medical Center for their dedicated patient care and clinical support. This work was funded by the American Academy of Neurology Resident Research Scholarship (to R.K.), the Simons Foundation’s SCGB program (to U.R.), and the BRAIN initiative through NIH grants U01NS117839 and R61DC022835 (to U.R. and A.N.M), and Columbia Engineering through start-up funds, the Dean’s Discretionary Research Funds, and the Maa-Liao Assistant Professorship of Biomedical Engineering (to N.R.). We also thank Jorge Aldana for assistance with computing resources.

## Author contributions

N.R. and T.G.A. conceived the study. N.R., U.R., T.G.A., A.P. and R.R. designed the behavioral task. R.K. and T.G.A. collected the data. R.K. and T.G.A. performed data processing. T.G.A. and R.R. performed data analysis. T.G.A., N.R., and U.R. wrote the initial draft of the manuscript. R.R. and A.P. designed and trained the vision-language model. R.R., R.K., A.N.M., A.P., N.R., and U.R. reviewed and edited the manuscript. C.M.R. provided patient care and facilitated experiments. A.N.M. performed all surgical procedures. U.R. and N.R. acquired funding.

## Methods

### Participants

Eleven adult patients (6 female; age range: 25–55 years; see Extended Data Table 2) with drug-resistant epilepsy participated in this study. All patients were undergoing stereotactic depth electrode implantation for seizure localization and potential surgical treatment at Cedars-Sinai Medical Center. Electrode placement was determined solely on clinical grounds by the clinical care team. Each patient provided written informed consent [46] to participate in research protocols approved by the Institutional Review Board of Cedars-Sinai Medical Center.

### Verbal instruction task

Patients performed a vision-language decision-making task in which they read written instructions for 2.5 s and viewed a sequence of two images for 1 s each. Patients then made a binary choice within 3 s, related to the contents of the images as determined by the trial’s instruction (Fig. 1a). Stimuli could belong to one of four categories: faces, cars, animals, and fruit. Instructions could be one of four task types: comparing the age of faces/cars, comparing the color of cars/animals, comparing the item count in animals/fruit, or comparing the identity of faces/fruit. Patients performed the task by responding to the instruction in the response phase, in which they were prompted to perform a binary button press. In age, color, and count tasks, patients had to answer “first” or “second” to report which of the two stimuli was the correct answer to the instruction prompt (e.g., “choose the oldest car”). In identity trials, patients had to answer “yes” or “no” to the identity comparison question (e.g., “are these two images of the same person?”). Each stimulus was defined as low/high along each of its feature axes (age, color, count) in relation to the other stimulus in its pair by human visual inspection. These binary split labels were used in all of the stimulus feature analyses. The feature along one axis was independent of the feature along the other axis. The left–right position of the response labels (“first” and “second”, or “yes” and “no”) was randomized across trials, so that patients could not plan the motor response before the response phase appeared. This dissociated decision formation from motor planning during stimulus presentation. Each patient performed 192 trials, counterbalanced to have the same number of trials for each stimulus category and for each task type. Trials were distributed across four blocks of 48 trials, with self-timed breaks in between. In half of the trials, instruction framing was a positive instruction (e.g., “choose the oldest car”; *pro-task* ), while the remaining half was a negative instruction (e.g., “do not choose the oldest car”; *anti-task* ). Identity trials had no framing and were thus excluded for framing-related analyses. Additionally, to probe the effects of instruction availability on visual stimulus processing, in half of the trials instructions were presented before stimulus presentation (*early instruction*), while in the remaining half they were presented after the first stimulus (*late instruction*), ensuring that a portion of stimuli were viewed without task context. Before the beginning of each trial, patients fixated on a central fixation cross for 1 s.

### Vision Language Model (VLM)

To model the computations required for this task, we built a multimodal vision-language model (VLM) consisting of separate vision and language encoders whose outputs were passed to a decision recurrent neural network (RNN; Fig. 1b). On each trial, the VLM received a natural language instruction with the same components as those given to patients, along with two RGB stimulus images. The RNN was required to produce a response corresponding to the image that satisfied the instruction. We trained 5 VLM instances with different random initialization seeds, each on a distinct image set from the one experienced by patients.

Specifically, the text encoder was a Sentence-BERT model (sbert big, sentence-transformers/bert-large-nli-mean-tokens) [47]. Images were encoded using CLIP ViT-B/32 (openai/clip-vit-base-patch32) [48], with the HuggingFace CLIPVisionModelWithProjection and CLIPProcessor classes used for feature extraction and preprocessing.

Both the text encoder and the vision encoder produced high-dimensional pretrained representations whose dimensionalities depended on the backbone architecture (e.g., 1024 for BERT-large and 512 for CLIP ViT-B/32). To align both modalities into a shared representation space, we introduced learned linear projection heads.

Let **h**_text_ ∈ R*^d^*^text^ denote the pooled instruction embedding produced by the Sentence-BERT encoder, and let **h**_vis1_, **h**_vis2_ ∈ R*^d^*^vis^ denote the CLIP image embeddings for the two candidate images. Each modality was first mapped into a shared 512-dimensional space via learned affine projections:

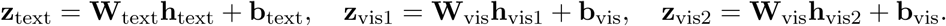

The decision network received a time-varying input that mirrored the time course of the experimental trial. Each trial was simulated as a sequence of steps divided into epochs: fixation, instruction, first stimulus, second stimulus, response-label cue and response. The three middle epochs were ordered as instruction–stimulus 1–stimulus 2 on early-instruction trials and stimulus 1–instruction–stimulus 2 on late-instruction trials, matching the trial condition experienced by the patient. The input at step *t* was the concatenation

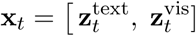

where each component was non-zero only while the corresponding information was on the screen: 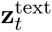 equaled the projected Sentence-BERT embedding of the instruction during the instruction epoch and 0 otherwise; 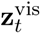 equaled the projected CLIP embedding of the first image during the first-stimulus epoch, that of the second image during the second-stimulus epoch, and 0 otherwise. During fixation and throughout the response epoch the input was entirely zero, so the network had to carry the instruction, the stimuli and its decision in its recurrent state. In particular, on late-instruction trials the instruction entered the network only after the first stimulus had been presented, exactly as for the patient.

The decision network was a gated recurrent unit (GRU) with 256 hidden units that processed **x***_t_*to produce hidden states **h***_t_*and a scalar output *y*^*_t_*:

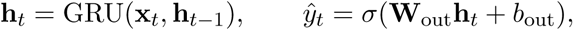

with **h**_0_ = 0 at the start of every trial. The response was read out as the mean of *y*^*_t_* over the final 10 steps, and a trial was scored as answered only if the final mean lay above 0.8 or below 0.2.

While the lower layers of both Sentence-BERT and CLIP were frozen to preserve their general semantic and visual representations, the final transformer blocks were fine-tuned jointly with the projection layers and decision module. Instruction is withheld during the first stimulus of late trials and the hidden state is reset in new trials. The sBERT regressor for the first stimulus of late trials is not passed forward in the network. The model is set up to have 500 steps along the following periods: stim1 period (early instruction) = [149,224]; stim1 period (late instruction) = [49,124]; stim2 period = [225,300]; instruction period (early instruction) = [49,149]; instruction period (late instruction) = [124,224]; response period = [349,500]. The model’s optimizer was AdamW, under three parameter groups with separate optimizers and learning rates: decision RNN 1e-4, vision encoder 5e-6, text encoder 1e-6; weight decay 0 for all three. Exponential learning-rate decay was *γ* = 0.99 for all three, stepped every 5,000 batches. Gradient-norm clipping was 1.0. Dropout was p = 0.05 on the embeddings.

### Visual text embeddings

The aforementioned VLM analysis is in part designed to provide regressors that test whether individual neurons correlate with instruction semantics via text embeddings. Assuming a subset of neurons do correlate with VLM text embeddings, we must still rule out the possibility that neural activations are better explained by the visual aspects of text (e.g. the geometric shapes of letters and words on the screen) than the semantic content of the instructions. To enable this comparison, we obtained 500x500 pixel screenshots of every instruction screen viewed by patients and obtained their visual embedded representation (i.e. not semantic) with a vision encoder (Fig. 2f). Specifically, we provided screenshots to a pretrained CLIP (one seed, ViT-B/32 architecture) model with no fine-tuning, which encodes images via a patch-based vision transformer followed by a projection head to produce a 512-dimensional embedding [48]. We then stored these embeddings for each trial to use as controls in the single-neuron VLM alignment analysis (see *Predicting neural activity from VLM* ).

### Decoding task components from the RNN

In Figure 2i, we decoded color, count, age, task, and target feature from the decision module of the VLM (the sensorimotor RNN). To do so, we performed a linear discriminant analysis (LDA) on the state of the RNN during the stimulus presentation period, stacking stimulus 1 and stimulus 2 as separate examples in the same vector to be decoded. We performed decoding separately for each of the 11 task sessions, measuring the RNN activations given the same inputs seen by patients, and each of the 5 network seeds, resulting in 55 runs, and averaging RNN activity during the stimulus presentation window. Label values marking “not applicable” (e.g. identity trials for target stimulus; stimuli whose category lacks the relevant feature axis) were excluded. Within each run, a linear classifier (variance filter, *z*-scoring, PCA to 20 components, and LDA with Ledoit–Wolf shrinkage) was evaluated with 5-fold stratified cross-validation and scored by balanced accuracy, so that chance was 1*/k* for *k* classes regardless of class imbalance. Within a session, the stimulus features are fully determined by the image pair, so cross-validation folds were therefore grouped by image pair (the sorted pair of image identifiers, 24 possible pairs per session), which places all trials using a given pair on the same side of every split. This prevents the classifier from scoring above chance by recognizing a pair or a trial it had seen in training. For each run and variable, a null distribution was obtained from 50 label permutations passed through the identical pipeline with the cross-validation partition held fixed. Scores and null scores were then averaged across the 5 seeds within each session, giving one value per session (*n* = 11) as the statistical unit. Significance was assessed with a permutation test, by obtaining a one-sample *t*-statistic across sessions against the empirical chance level (the mean of the null), with the null distribution of the statistic generated by 10,000 bootstrap draws that sampled one permutation per session; *P* -values were corrected across epoch windows within each variable by the Benjamini–Hochberg procedure.

### Behavioral analysis

All behavioral analyses (for both patients and models) were aggregated across sessions (or across initialization seeds for models) for inferential statistics (Fig. 2). Trial-level accuracy was defined as a binary variable indicating whether the response matched the correct answer on that trial, where each trial had an unambiguously defined correct answer given the instruction. Missed trials due to response timeouts were considered incorrect.

Group-level statistical analyses were conducted across sessions using paired nonparametric tests. Specifically, two-sided paired Wilcoxon signed-rank tests were used to compare condition-specific accuracy measures at the session level, including comparisons between pro-task and anti-task trials, early-instruction and late-instruction trials, and all pairwise comparisons across stimulus categories and task types. For comparisons involving more than one pairwise test (e.g., the four-level category and task-type analyses), raw *P* -values were corrected for multiple comparisons using the Benjamini–Hochberg false discovery rate (FDR) procedure at *α* = 0.05.

### Neural recordings and preprocessing

The neural recording and preprocessing pipelines used in this study were the same as those described in a companion study [14], and the details on recording and spike sorting are reproduced here. Intracranial recordings were obtained at Cedars-Sinai Medical Center from bilateral hybrid depth electrodes, each containing eight microwires (AdTech Medical). Electrode implantation sites were determined exclusively by clinical needs for seizure localization, and included the ventral temporal cortex (VTC), dorsal anterior cingulate cortex (dACC), pre-supplementary motor area (preSMA), ventromedial prefrontal cortex (vmPFC), hippocampus (HPC), and amygdala (AMY) (Extended Data Table 2). Broadband extracellular signals (0.1–9,000 Hz) were continuously recorded from each microwire at a sampling rate of 32 kHz using the ATLAS system (Neuralynx Inc.). Recordings were locally referenced within each electrode bundle to a designated low-impedance microwire.

Electrode localization was performed by co-registering postoperative computed tomography (CT) scans with preoperative magnetic resonance imaging (MRI) using standard alignment procedures implemented in FreeSurfer (v5.3.0 and v7.4.1) and Advanced Normalization Tools (ANTs v2.1) [49]. Individual preoperative MRIs were aligned to the MNI152-space CIT168 probabilistic atlas for visualization of electrode locations. All electrode localization was performed in the participant’s native space, and electrode contacts were confirmed to be within gray matter through visual inspection.

Spike detection and sorting were performed offline using the semi-automated template-matching algorithm OSort (v4.1 [50]). The raw broadband signal was band-pass filtered between 300 and 3,000 Hz, and candidate spikes were detected using an adaptive threshold. Putative single units were isolated based on waveform shape, refractory-period violations, and cluster separation. Units that did not meet these isolation criteria were excluded. Only patients with at least one well-isolated single neuron in one of the regions of interest were included in the analyses.

### Encoding analysis

To quantify the proportion of neurons tuned to each task variable, we fit a single generalized linear encoding model per neuron. The response variable was the firing rate in the 1-s stimulus presentation window, referenced to stimulus onset. First and second stimulus presentations were treated as separate observations and concatenated, yielding 192 × 2 = 384 observations per neuron. All presentations entered the model (correct and incorrect trials, all four task types).

The design matrix contained main effects for ten task variables, grouped as bottom-up, top-down and task-state factors. Bottom-up factors were the stimulus category (face, car, animal, fruit; three indicator regressors) and the continuous age, color and count values of the presented stimulus, each *z*-scored across the presentations for which the axis applies to the stimulus category and set to zero otherwise. Top-down factors were the instructed task (age, color, count or identity comparison; three indicators), task framing (coded +1 for pro-task instructions, −1 for anti-task instructions and 0 for identity-matching trials, whose instructions carry no framing), the target feature and the correct choice. The target feature is whether the presented stimulus was the high or the low item of its pair on the instructed axis (one indicator, high = 1); the correct choice is whether the first or the second stimulus of the trial was the target (one indicator, second = 1). To control for trial difficulty, we added reaction time in the decision phase as a regressor of no interest in the model, as a proxy for the difficulty of each trial. We observed similar results without the inclusion of reaction time as a regressor.

Both target feature and correct choice variables exist only when the trial is a feature-comparison trial (i.e. not a match trial) and the instruction has already been delivered: on identity-matching trials there is no feature axis and no target, and before the instruction the patient cannot know which feature is relevant or which stimulus will be the target. The presentations on which the two variables are undefined are therefore the same set, namely both presentations of every identity trial and the first stimulus of every late-instruction trial. On these presentations both indicators were set to 0 and a single additional regressor, *undefined*, was set to 1 (0 elsewhere). This regressor is not a task variable and is never tested; its role is to give the undefined presentations their own mean in the model, so that they are not treated as “low” and “first stimulus” by default when the remaining variables are tested. Task-state factors were the position of the stimulus in the sequence (first or second) and the instruction order (early or late).

Since spike counts are non-negative, over-dispersed and frequently zero, the rate *y* was modeled with the same Tweedie generalized linear model used for the model-neuron alignment (see *Predicting neural activity from VLM* ): a log link, E[*y*] = *µ* = exp(*Xβ*), and a power variance function Var(*y*) = *ϕ µ^p^* with *p* = 1.1, which places the error distribution between the Poisson (*p* = 1) and gamma (*p* = 2) families and accommodates excess zeros [51]. The model

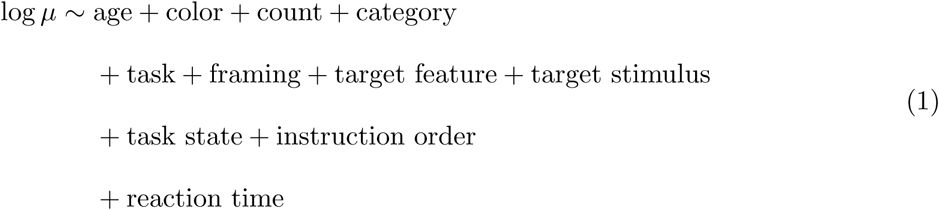

was fit by maximum likelihood (iteratively reweighted least squares) without regularization.

Tuning to each variable was assessed by an analysis-of-deviance *F* -test [52]: the full model was compared with the model lacking that variable’s regressors,

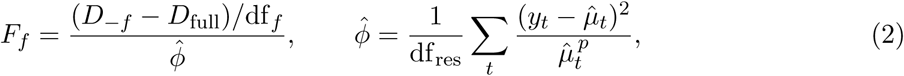

where *D* denotes the Tweedie deviance of the indicated model, df*_f_* the number of regressors of variable *f* , and *ϕ*^^^ the Pearson estimate of the dispersion from the full model with df_res_ residual degrees of freedom; *F_f_* was referred to an *F* (df*_f_ ,* df_res_) distribution. Each variable’s test is therefore conditional on all others. The tests of the eight variables that are defined on every presentation (category, the three feature values, task, framing, task state and instruction order) used all 384 presentations. The tests of the target feature and of the target stimulus were run on the presentations on which they are defined (feature-comparison trials after the instruction): the undefined presentations were excluded, the *undefined* regressor was dropped (it is constant on the remaining rows), and the full and reduced models were fit on this subset with all other regressors retained. Each of these two tests is therefore a contrast between the variable’s two defined levels (high versus low on the instructed axis; first versus second stimulus as target), conditional on the other variables, and cannot be driven by differences between defined and undefined presentations, such as whether the rule was known or whether the trial was an identity trial. A neuron was considered tuned to a variable at *P <* 0.05. The proportion of tuned neurons in each brain area was compared with the 5% false-positive rate using a one-sided binomial test with sample size equal to the number of recorded neurons in that area.

### Top-down index

To summarize each neuron’s role in encoding sensory versus task-related factors, we defined a top-down index (TDI), a normalized measure of how much unique variance in a neuron’s firing is explained by bottom-up stimulus variables relative to top-down task and goal variables. For each session, we assembled a 14-column design matrix *X* ∈ R*^N×^*^14^, with *N* = 192 × 2 = 384 examples, in which columns are six stimulus-related factors *S*, seven task-related factors *T* , and one indicator *z* for correct versus incorrect response. Specifically, *S* contained color, count, and age features, as well as one-hot coded stimulus categories (3 dummy variables since 4 categories imply 3 degrees of freedom). *T* contained one-hot coded task type (4 dummy variables, since there were 4 task types + 1 undefined type for late instruction trials, totaling 4 degrees of freedom), task framing, target stimulus, and target feature. We added the indicator for correct versus incorrect response to isolate variance related to errors, and subsequently correlate the resulting TDI in different brain areas with error rates. Neural responses in the stimulus presentation window (treating first and second stimuli as separate examples) were correlated with the design matrix *X* composed of a concatenation of *S* and *T* , as well as the error regressor *z*.

We then proceeded to estimate the unique partial contributions of *S* and *T* by using three nested ordinary least squares (OLS) regressions: a full model including all regressors (*y* ∼ [*S, T, z*]); a model dropping the stimulus block (*y* ∼ [*T, z*]), and a model dropping the task block (*y* ∼ [*S, z*]). Model predictions were computed via OLS (with scikit-learn LinearRegression), and fits were summarized by sum of squared errors (SSE).

The unique variance explained by stimulus variables, conditional on task variables and correctness, was computed as a partial *R*^2^ quantity:

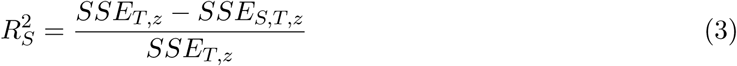

Similarly, unique variance explained by task variables conditional on stimulus variables and correctness was computed analogously:

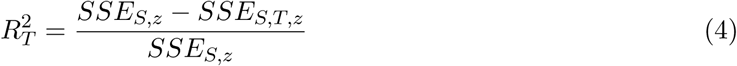

Finally, the top-down index (*TDI* ∈ [−1, 1]) was defined for each neuron as the normalized difference between these unique variance components:

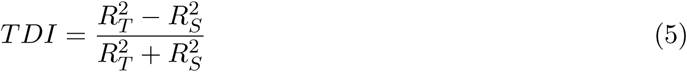

In this definition, positive values indicate relatively stronger unique contributions of top-down or task factors, while negative values indicate relatively stronger unique contributions of bottom-up or stimulus factors.

### Decoding analysis

We performed neural decoding by using pseudopopulations of firing rate matrices within the stimulus presentation window, making the simplifying assumption that all brains were recorded at the same time. We counted spikes in 300 ms windows with a 25-ms sliding step and kept neurons whose average firing rate in the session was above 0.5 Hz, referencing time bins to the window starts. This resulted in a total *n* = 743 neurons, 148 in VTC, 243 in MTL, 118 in vmPFC and 234 in MFC. After counting spikes within each time bin, we performed temporal smoothing via 1D convolution with a causal exponentially decaying kernel. Concretely, let *r*(*t*) denote the raw firing rate at discrete time bin *t*, obtained from sliding-window spike counts with step Δ*t* = 25 ms. Temporal smoothing is performed by convolving the rate sequence with a kernel *h*(*k*):

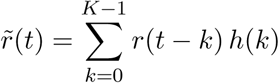

where *r̃*(*t*) is the smoothed firing rate and *K* is the kernel length.

The kernel is defined as a delayed exponential:

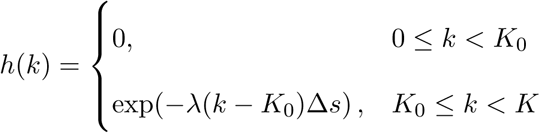

where

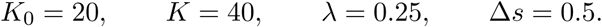

To assemble pseudopopulations for each variable of interest, we constructed a condition-balanced list of trial indices for the possible levels of the variable. Each trial in the pseudopopulation matrix is formed by taking one random trial that matches a particular combination of desired task variables and concatenating neurons across sessions. Since alignment across sessions is arbitrary and trial selection within condition is random, we perform this sampling procedure 25 times, generating 25 possible pseudopopulation samples for each decoded variable. First, the target number of trials to assemble is set as 384, corresponding to the number of individual stimuli shown in a session. Out of these, we perform a 288/96 trial train/test split. We also bootstrapped a null distribution for decoding by randomly shuffling neural firing rates across trials 500 times for each of these 25 pseudopopulation resamples.

Within each train/test split, the number of pseudotrials allocated to each level of a variable of interest was determined by dividing the total number of train/test trials by the number of unique conditions. For each condition, we identified all trials in each session belonging to that condition and randomly sampled the required number of indices. When the number of available trials to select from exceeded the required quota, trials were sampled without replacement. When fewer trials were available than required, we filled the trial quota by repeatedly sampling from the available trials across multiple rounds, effectively bootstrapping the condition so that each condition contributed the same number of pseudotrials. All decoded variables were binary (age, count, color, target feature, choice) with the exception of category, which had four possible values (cars, fruit, animals, faces).

Once train and test pseudopopulations were assembled, we performed classification with a linear SVM (StandardScaler() + SVC(kernel=’linear’, C=100)), evaluated independently at each time bin. Final decoding accuracy scores are reported corresponding to mean result across the 25 pseudopopulation resamples. The same averaging procedure was performed in the bootstrapped null distribution, and *P* -values were computed by comparing true decoding accuracy to bootstrapped shuffled decoding accuracies, up to *P <* 0.002. *P* -values were cluster corrected across time bins [53].

### Generalization analysis

The generalization analysis utilizes the same backbone as the decoding analysis (linear SVM, 25 pseudopopulation resamples), changing which trials belong to the train and test sets. Specifically, we trained decoders on pseudopopulation activity from one subset of trials defined by a generalization variable of interest and evaluated that decoder on a disjoint subset of trials defined by different values of the same generalization variable. We then utilized a generalization index to understand cross-condition generalization accuracy (CCGP), with within-condition decoding accuracy (*D*) as a baseline [17]:

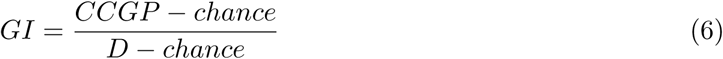

Chance levels were 0.25 for the category variable and 0.5 for the other tested variables. We determine GI significance periods by bootstrapping a null distribution with 500 permutations shuffling decoded labels and counting occurrences where true GI was larger than shuffled values to obtain *P* -values. Subsequently, we applied cluster correction across time bins [53].

### Single-trial populational representations

We developed a method for predicting how single-trial representations of stimuli are altered in the presence (or absence) of specific instructions. For this analysis we restricted the trial space to correct trials only. For each session *s* we constructed a matrix *X_s_* ∈ R*^nExamples×nNeurons^* by iterating over neurons and filling each column with the per trial spike counts in the stimulus presentation windows (384 examples, minus incorrect trials).

Then, we trained a neural network to predict neural activities contained in *X_s_* from the (instruction, stimulus) pair for each trial, with an 80*/*20 train-test split. To train this neural network, we designed a pipeline to represent instructions and stimuli in trial-wise learnable embeddings, allowing the embedding of instructions to gate the representation of stimuli.

First, the model embedded stimuli identities on a per-session basis, producing an embedding *E_s_* of dimension *d* = 32. Then, this session-wise representation was projected onto a latent representation *z* in a shared space across sessions (*d* = 64), via a multilayer perceptron (MLP) with ReLU activation and dropout (*p* = 0.1). Second, each verbal instruction is reduced to its fundamental one-hot-coded flags (instruction framing and comparison type, target feature), which are represented as a learnable context embedding *c* of dimension 64. At this stage, dropout (*p* = 0.1) is applied to the context embedding.

Then, we allow the context vector *c* to determine how the stimulus representation *z* should be transformed. To do this, we base our procedure on feature-wise linear modulation (FiLM) [54], in which a feature-wise affine transformation conditioned on context gates internal network representations. Briefly, FiLM works by adding a simple feature-wise linear transformation to a network’s internal representations, conditioned on a context vector. In our case, we allow the context vector *c* to generate 2 linear parameters which will transform the stimulus representation: bias and rate parameters *β* and *γ*, with *γ, β* ∈ R^64^. This is achieved with a two-layer MLP with learnable *W* and *b* parameters and dropout *p* = 0.1:

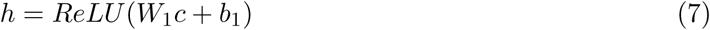

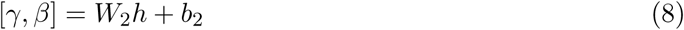

Finally, the FiLM rate and bias parameters modulate how stimulus representations are expected to behave under the presence of the context vector *c*:

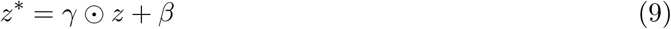

where ⊙ denotes element-wise multiplication. Essentially, *z* encodes what the stimulus is, *c* encodes what rule should be applied, and FiLM computes how the instruction rule transforms the stimulus representation.

To return to neural space and represent spike counts *y*, the model uses session-specific learnable linear decoders. For session *s*, the predicted firing rate vector for neurons recorded in that session is given by:

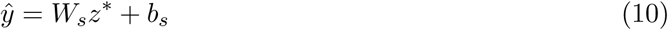

The loss objective to train this entire representation network, backpropagated with PyTorch, is obtained by comparing actual and estimated spike counts for each session:

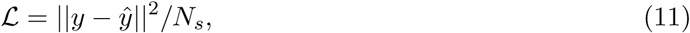

where y are the actual neural spike counts and *N_s_* is the number of neurons in session *s*.

After training the representation predictor, we quantified whether the model genuinely used instruction context and stimulus identity to generate neural predictions by constructing two null procedures: a context (instruction) null and a stimulus-identity null. Both tests operated exclusively on held-out test trials and compared the prediction error obtained under the correct variables to prediction errors obtained when one component of the input was systematically mismatched while keeping the other components fixed.

For the instruction-null test, the goal was to determine whether the predicted neural representation depended on the instruction context rather than on stimulus identity alone. For each test trial *i* in session *s*, the model first generated a neural prediction *y*^*^true^* using the correct stimulus identity and the correct instruction flags. The prediction error was computed as the mean-squared difference between this prediction and the observed neural firing-rate vector *y_i_*.

To generate the instruction-null distribution, the procedure held the stimulus identity fixed and systematically replaced the instruction flags with alternative context combinations (*n* = 500) observed in the same session. For each null context *c^′^ ≠ c_i_*, the model produced a new prediction 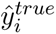 using the same stimulus but the substituted context flags. With this, we produced a set of null losses for that trial. These null losses quantified how well the model could predict the neural response when the stimulus was correct but the task instruction was incorrect. The mean of these null losses defined the instruction-null baseline for that trial. Across trials, we compared the vector of true losses to the vector of instruction-null losses using a paired Wilcoxon signed-rank test (one-sided, testing whether true losses were smaller). A significant result indicates that the correct instruction context improves prediction accuracy, implying that the model learned context-dependent transformations of stimulus representations.

For the stimulus-identity null test, the goal was to determine whether the model’s predicted representation corresponded to the correct stimulus identity rather than simply reflecting the instruction context. Similarly to the instruction-null case, we compare true firing rates with predicted estimates from shuffled stimuli. Specifically, to construct the stimulus null distribution, the procedure held the instruction context fixed and compared the prediction to neural responses associated with other stimuli presented under the same context within the same session. To enable this, the algorithm identified all other trials whose context flags matched the current trial’s context but whose stimulus identity differed. For each such trial we computed the mean-squared difference between the predicted neural vector and the actual neural vector observed for that alternative stimulus. This produced a set of stimulus-null losses that quantified how well the model’s prediction for the current stimulus matched neural responses evoked by different stimuli under the same task rule.

Together, these two null procedures isolate the contributions of the two input factors that define the model’s computation. The instruction-null test evaluates whether neural predictions depend on the correct task rule when the stimulus is fixed, while the stimulus-null test evaluates whether predictions depend on the correct stimulus when the task rule is fixed. Demonstrating lower prediction error under the true configuration in both tests indicates that the learned representation jointly encodes stimulus identity and task instruction and that the FiLM-mediated transformation of stimulus embeddings captures rule-dependent changes in neural population activity.

### Predicting neural activity from the VLM

We developed a pipeline to align human single-neuron activity to unit activation in the VLM, with the goal of explaining variance in human neurons and predicting their activity out-of-sample. This approach mirrors previous work that obtained computational insight into visual ventral stream activity by aligning it to convolutional neural network models of machine vision [19]. We performed alignment by treating each recorded neuron as an encoding target and asking how well activity in a specific module of the VLM (vision, language, or decision RNN) predicts that neuron’s firing rate in the stimulus presentation window. After the VLMs were trained (5 random initializations), we showed each network exactly the same sequence of trials experienced by the patient in each session and averaged the activity of each artificial unit within the stimulus presentation window (for the vision and language modules, the CLIP embedding of the presented image and the SBERT embedding of the trial’s instruction, respectively; for the RNN module, the 256-dimensional hidden state of the decision GRU). Each trial contributed two observations, one per stimulus presentation, so that a session with 192 trials yielded 384 paired samples (*X_t_, y_t,i_*), where *X_t_* is the model activation at presentation *t* and *y_t,i_* the firing rate of neuron *i* in the 1 s following stimulus onset.

To quantify alignment strength between each neuron and the model feature space, we fit a regularized Poisson–Gamma (Tweedie) GLM, appropriate for non-negative spike-rate targets. Prior to model fitting, we standardized the feature distribution with quantile mapping and reduced dimensionality with PCA to concentrate predictive variance into a 20-dimensional subspace that remained comparable across sessions and model seeds. The L2 regularization strength of the GLM was searched over the logarithmic grid np.logspace(1,-4,5) by nested cross-validation (an inner 4-fold loop to select the hyperparameter, nested within an outer 4-fold loop to obtain held-out performance). Concretely, we used the following sklearn pipeline to regress neural firing rates on artificial unit activations:

pipe = Pipeline(

[

("scaling", QuantileTransformer(n_quantiles=80)), ("pca", PCA(n_components=20)),

("regress", linear_model.TweedieRegressor(power=1.1, warm_start=True, max_iter=200)),

]

)

Presentations were assigned once, by a seeded random draw, to the four outer folds, and the identical assignment was used for every neuron, module, network seed and permutation, so that all model comparisons described below were evaluated on the same held-out presentations. We quantified alignment as the out-of-sample fraction of deviance explained,

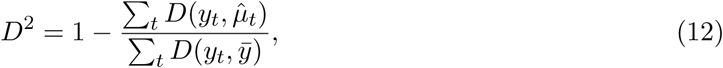

where *D*(*y, µ*) is the Tweedie unit deviance (Eq. 13), *µ*^*_t_* the prediction for held-out presentation *t* from a model fit on the remaining folds, and *ȳ* the mean rate of the training presentations. *D*^2^ is the deviance-based analogue of *R*^2^ for a GLM (both equal the fraction of null deviance explained; for a Gaussian model they coincide) and is the natural goodness-of-fit measure for Tweedie regression [51]. For each outer fold we trained the full preprocessing and regression pipeline on the training presentations and evaluated it on held-out presentations, producing one *D*^2^ value per fold, which we averaged to obtain a single generalization metric per neuron. This ensures that variance explained pertains to unseen presentations, above and beyond goodness-of-fit on the trials used for parameter estimation. Out-of-sample predictions and fold assignments were stored for every neuron and reused by all downstream analyses.

To determine which neurons showed out-of-sample alignment beyond what was expected by chance, we built a null distribution of *D*^2^ for each neuron by permuting the order of trials (keeping the two presentations of a trial together) and repeating the fitting procedure (*n* = 500), with the regularization strength fixed to the values selected on the unpermuted data. For each neuron we defined a significance threshold as the 99th percentile of its null *D*^2^ distribution and labeled the neuron *aligned* when its real *D*^2^ exceeded this neuron-specific cutoff. To determine whether the number of aligned neurons in each brain area was greater than expected by chance, we used the same one-sided binomial test as in the single-neuron encoding analysis. Alignment was computed separately for each of the five trained networks; we report results for the seed-averaged *D*^2^.

### Post-hoc characterization of VLM-aligned neurons

A neuron can be predictable from the model for various reasons: the model’s activations carry the task’s structure (which feature was instructed, whether the instruction had already been delivered, the stimulus category), so any neuron that encodes these variables will be partly predictable from any module that represents them. To determine what the alignment reflects, we subjected every aligned neuron to a battery of nested model comparisons that ask (i) whether the neuron’s tuning is accounted for by the task variables of the single-neuron encoding model and, if so, by which ones, and (ii) whether the VLM module predicts variance in the neuron that these task variables cannot. The battery was run separately for each module (vision, language, RNN); all tests below were computed on the same 384 presentations and, where applicable, the same four folds as the alignment, and no model was refit for the alignment statistic itself.

#### Linear task-variable design

We used the same main-effects design as in the single-neuron encoding analysis (Methods, *Single-neuron encoding*): the presented stimulus’s continuous value on the age, color and count axes (*z*-scored within the categories to which the axis applies and set to zero where the axis is undefined), stimulus category, instructed feature, task framing (a single signed column, +1 pro, −1 anti, 0 identity), whether the presented stimulus was the correct target (first or second stimulus, defined for non-identity trials once the rule was known), the presented stimulus’s within-pair rank on the instructed axis, task state (first versus second stimulus) and instruction order (early versus late). Categorical variables were one-hot coded with an explicit “undefined” level where a variable was not defined for a presentation. The design contained main effects only; no interaction terms were included.

#### Per-factor tuning and explanatory labels

For each aligned neuron and each factor *f* we tested whether *f* explained variance beyond the remaining factors with a nested *F* -test between the full main-effects model and the model with *f* removed, fit by ordinary least squares to the square-root-transformed rate (the variance-stabilizing transform for Poisson-like counts, which brings the analytic *F* distribution close to its nominal calibration for sparse spike counts; see *Single-neuron encoding* ). The unique share of each factor was *L_f_* = (SS*_−f_* − SS_full_)*/*SS_null_, where SS denotes the residual sum of squares of the indicated model. A neuron was tuned to *f* if *P_f_ <* 0.05 and *L_f_ >* 0. We grouped factors into bottom-up (category, age, color, count), top-down (instructed feature, framing, target stimulus, instructed-axis rank) and task-state (first/second stimulus, instruction order) variables. Each neuron’s primary task-variable label was the significant factor with the largest unique share *L_f_* ; neurons with no significant factor received no task-variable label.

#### VLM content beyond the task variables

To ask whether a module predicts variance that the task variables cannot, we tested the increment of the module over the design with the same nested *F* -test: the 20 principal components of the module activations used by the encoder (standardized) were added to the full main-effects design, and the *F* statistic for the added block (20 numerator degrees of freedom) was computed against the design-only model. A neuron was labeled *VLM beyond design* when *P <* 0.05 and the module increased the fraction of deviance explained. For the vision module, a beyond-design neuron is, by construction, tuned to the identity or continuous appearance of the presented item beyond its category and feature value (“visual item tuning”), and for the language module to the wording of the instruction beyond the instructed feature and framing (“instruction wording”). For the RNN, this corresponds to a task solution space which combines these visual and linguistic inputs to perform decisions.

#### Explanation of each aligned neuron

Each aligned neuron received one primary explanation, with precedence *VLM beyond design >* task-variable label *> unexplained* : neurons that passed the beyond-design test were labeled VLM beyond design; otherwise the neuron carried its primary task-variable label; and neurons that passed neither test were labeled *unexplained*. An unexplained neuron is thus one that is predictable from the VLM at better than chance level, but for which neither any single task variable (given the others) nor the VLM (given all task variables) reaches significance in the linear model. This occurs when weak tuning is distributed across several task variables, when the alignment is marginal, or when the neuron’s dependence on the task variables is not additive. We report, per brain area, the percentage of aligned neurons in each explanation category.

### Behavioral relevance of the alignment

Post hoc, to determine the behavioral relevance of the neural–model alignment, we tested whether predictive quality in aligned neurons differed between correct and incorrect trials. After fitting the Tweedie model, we generated predictions for every presentation and computed a Tweedie deviance per sample, which acts as a likelihood-based loss for Tweedie GLMs [51]. For a Tweedie generalized linear model with mean *µ* and variance function Var(*Y* ) = *ϕµ^p^*, where *y* denotes the observed response, *µ* the predicted mean, *p* the Tweedie power parameter, and *ϕ* the dispersion parameter, the *unit deviance* for *p* = 1.1 is

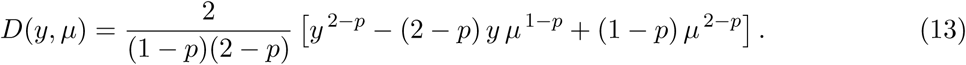

With this relationship between predictions and deviance, we obtained the average deviance across samples separately for correct and incorrect trials. For each neuron, we computed and report the difference in deviance between behaviorally correct and incorrect trial sets:

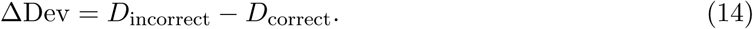

By this definition, positive ΔDev values indicate a worse fit (higher deviance) on incorrect than on correct trials. We restricted this analysis post hoc to the subset of neurons that passed the alignment significance test, so that ΔDev effects describe behavior-dependent changes in meaningful model–neuron alignment. At the group level, we summarized ΔDev by brain area using the mean and standard error of the mean across neurons and tested whether ΔDev in each brain area differed from zero with a one-sample two-sided t-test of delta Dev against zero across aligned neurons, Benjamini-Hochberg corrected across the four areas.

Regarding correlation with text embedding, we also controlled for the possibility that neurons are better explained by low-level visual representations than semantic text embeddings (i.e. the visual shape of letters and words on the screen). For this, we utilized 512-dimensional visual text embeddings based on screenshots of instruction panels experienced by patients (see *Methods*, *Visual text embeddings*, for more details on how we obtained such embeddings). We then correlated neural activity to these 512-dimensional embeddings in the same manner as the sBERT/CLIP embeddings obtained from the VLM. Post-hoc, we selected the subset of neurons found to be significantly explained by the sBERT text embeddings and performed a model comparison to test whether each one was better explained by semantic-based embeddings or visual-based embeddings. We performed this model comparison by directly comparing the cross-validated *D*^2^ metric for each neuron. We then obtained the difference in *D*^2^ deviance for each neuron, comparing the sBERT embedding GLMs versus the visual text embedding GLMs. We averaged these differences across VLM seeds and cross-validation folds, and performed a one sample, two-sided *t*-test over this *D*^2^ difference distribution, across neurons, to determine whether each brain area’s population was better explained by semantic sBERT embeddings or visual text embeddings.

### Expansion of the stimulus representation after the instruction

In the task solution space of the VLM, the population representation of a stimulus is compact before the instruction is known and expands, in an instruction-dependent manner, once the instruction is known (Fig. 5). We quantified this in the neural data and the RNN (Extended Data Fig. 9) by comparing the first stimulus of late-instruction trials (presented before any instruction) with the first stimulus of early-instruction trials (presented after the instruction): the same images, in the same serial position within the trial, differing only in whether an instruction preceded them. For each neuron we took the square root of the firing rate in the 1 s following stimulus onset and *z*-scored it over all presentations of the session.

Presentations were assigned to conditions defined by stimulus category × instructed feature × framing, using the same labels before and after the instruction; before the instruction, these labels denote the instruction the patient would receive later in the trial, so any condition structure along them at that time cannot reflect the instruction and serves as a built-in negative control.

Because the estimated centroid of a condition contains trial-to-trial noise, the naive variance of condition centroids is positively biased by an amount that depends on the noise level and the number of presentations per condition and would therefore differ between the pre- and post-instruction comparisons for reasons unrelated to representational geometry. We therefore used cross-validated estimates, in which each squared deviation is replaced by the product of the deviations estimated from two independent halves of the presentations; this estimator is unbiased, has an expectation of zero when conditions do not differ [55].

For this, within each session and condition, presentations were randomly split into two halves, and condition centroids were computed separately for each half. For every neuron we then computed the *instruction-related spread*, the variance of condition centroids around their stimulus category centroid, as the product of the half-A and half-B deviations summed over conditions (weighted by the number of presentations) and the *total centroid variance* (variance of all condition centroids around the grand mean). Per-neuron values were averaged across the random split (10 repetitions) and summed over the neurons of a brain area, pooled across sessions, and divided by the number of neurons so that areas are comparable irrespective of their size.

The instruction-related spread was compared before and after the instruction by its difference, and the expansion of the occupied volume by the difference in total centroid variance (after minus before). Confidence intervals were obtained by bootstrap over neurons (500 resamples). Significance of the before-after difference was assessed with a permutation test in which the early/late trial labels were shuffled within session and condition (200 permutations), which preserves the condition structure and the number of presentations while removing any dependence on whether the instruction had been delivered. The RNN values were obtained by applying the identical computation to the network: the 256 units of the decision RNN, averaged over the steps of the same stimulus epoch on the same trials (Methods, *Predicting neural activity from the VLM* ), were treated as a population, *z*-scored per unit, and the instruction-related spread and total centroid variance before and after the instruction were computed with the same split-half procedure and bootstrap over units, averaged over the five trained network instances.

## Code Availability

The code for the analyses performed in this work will be made publicly available upon acceptance.

## Data Availability

Data used in this study will be made publicly available upon acceptance.

---

**Extended Data Fig 1.**
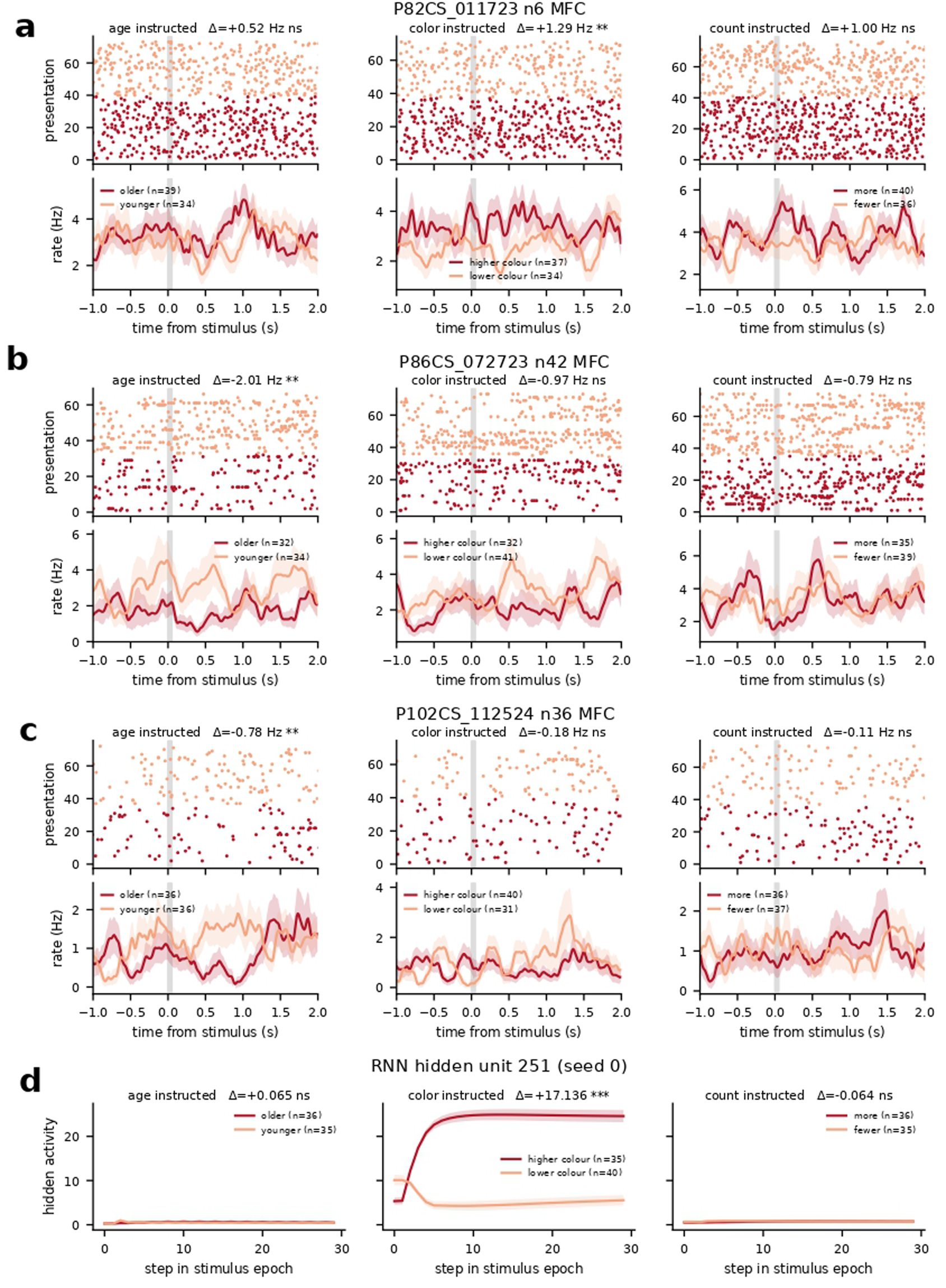
Target feature coding in MFC neurons and RNN. **a–c**, Raster plots of 3 MFC target feature neurons, firing specifically for low vs. high value of specific instructed features. **d**, RNN node that displays the same target feature axis specific code. Two-sided Mann-Whitney U test of the spike counts within stimulus presentation window, or mean RNN node activation, contrasting high vs low. The reported Δ is mean(high) - mean(low) in Hz.(***: *P <* 0.001, **: *P <* 0.01, *: *P <* 0.05).

**Extended Data Fig 2.**
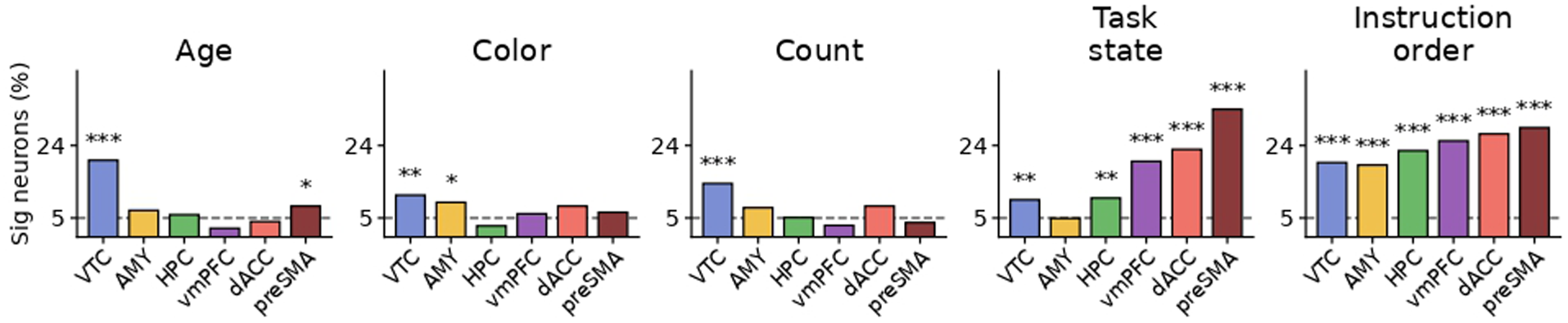
Single neuron correlations with extra factors in encoding analysis. **a–e**, Proportion of neurons correlated during stimulus presentation with stimulus age, color, count, current task state (first vs. second stimulus), and instruction order (***: *P <* 0.001, **: *P <* 0.01, *: *P <* 0.05, binomial test).

**Extended Data Fig 3.**
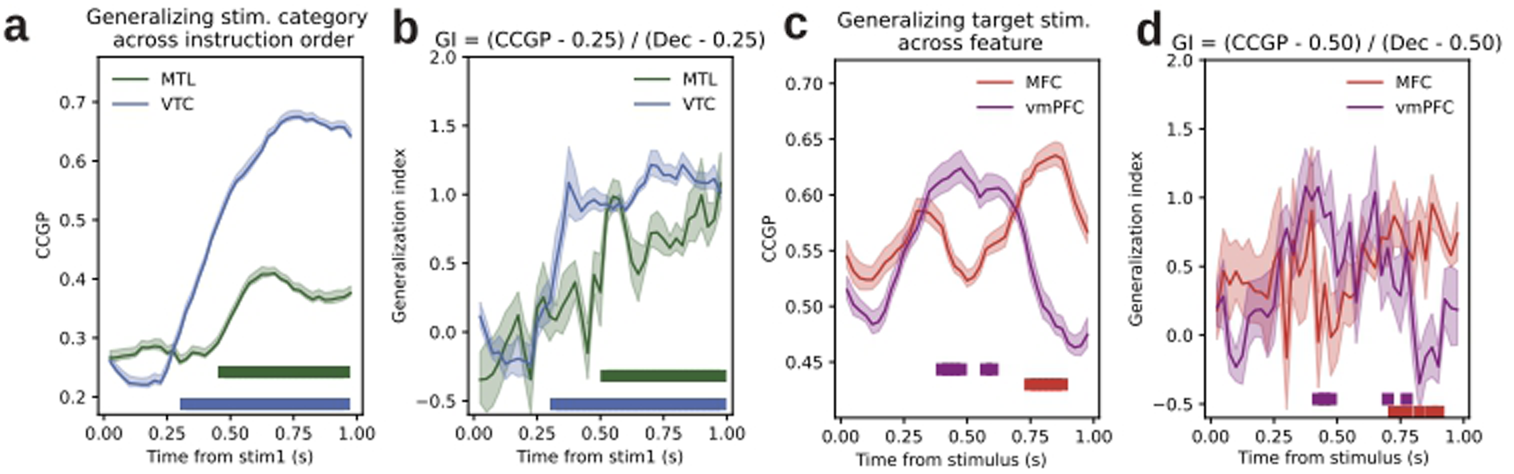
Generalization analyses inverting train/test sets with respect to main text. In all panels shaded areas indicate s.e.m. and bars indicate significant time bins (permutation test, cluster-corrected across time bins). **a**, CCGP for stimulus category across instruction order (early vs. late). Decoders were trained in late instruction trials and tested in early instruction trials. **b**, Generalization index for category across instruction order. Decoders were trained in late instruction trials and tested in early instruction trials. **c**, CCGP for target stimulus (first vs. second) across target feature magnitude (low vs. high). Decoders were trained in high feature level trials and tested in low feature level trials. **d**, Generalization index for target stimulus (choice) across target feature. Decoders were trained in high feature level trials and tested in low feature level trials.

**Extended Data Fig 4.**
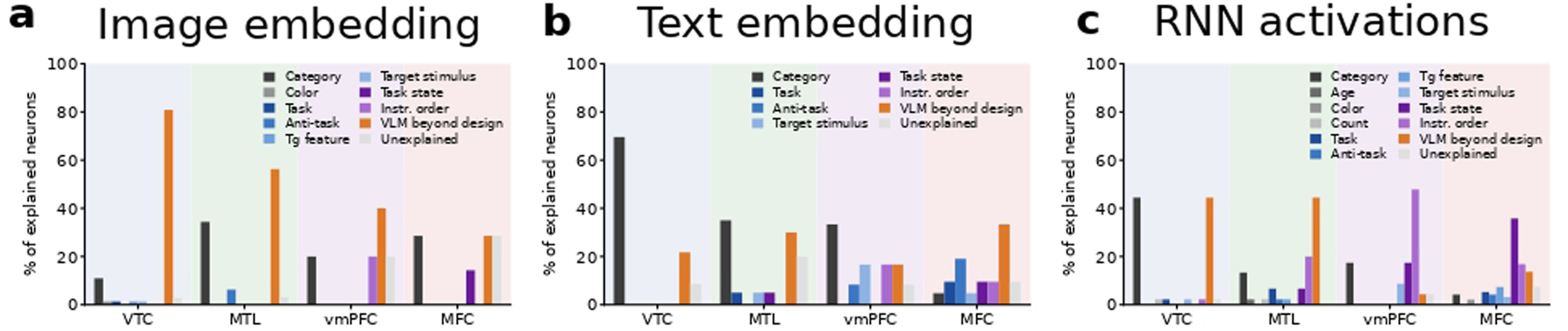
Explanatory factors for each neuron which correlates with the VLM. **a**, Proportion of neurons explained by the CLIP image embeddings which are better explained by each task factor, or better explained by the VLM. **b**, Same, for sBERT text embeddings. **c**, Same, for RNN activations.

**Extended Data Fig 5.**
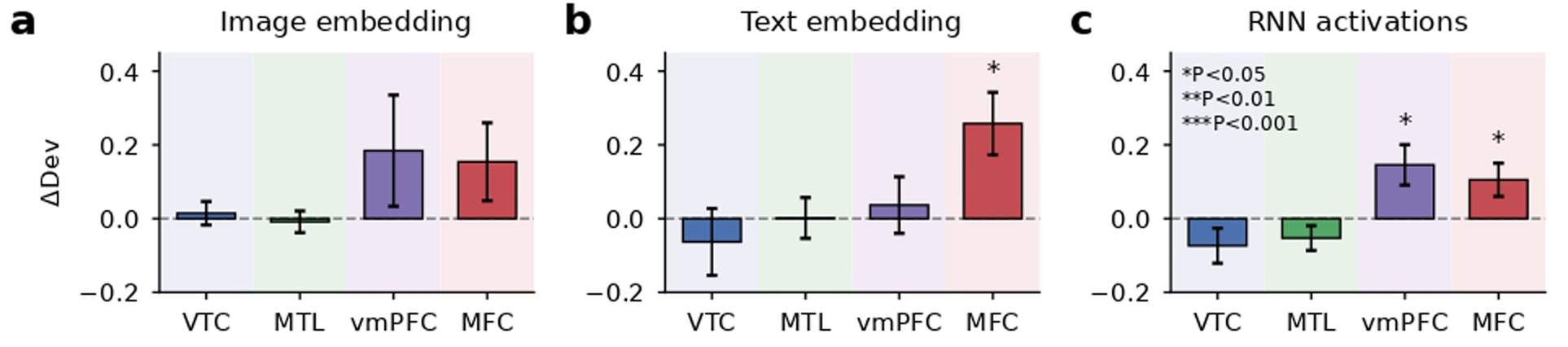
Neural-VLM alignment correlates with behavioral performance on the task. **a**, Change in model deviance in predicting neural activity in correct versus incorrect trials, from image embeddings (*: *P <* 0.05, One-sample two-sided t-test of delta Dev against zero across aligned neurons, Benjamini-Hochberg corrected across the four areas). **b**, Same, for text embeddings. **c**, Same, for RNN activations.

**Extended Data Fig 6.**
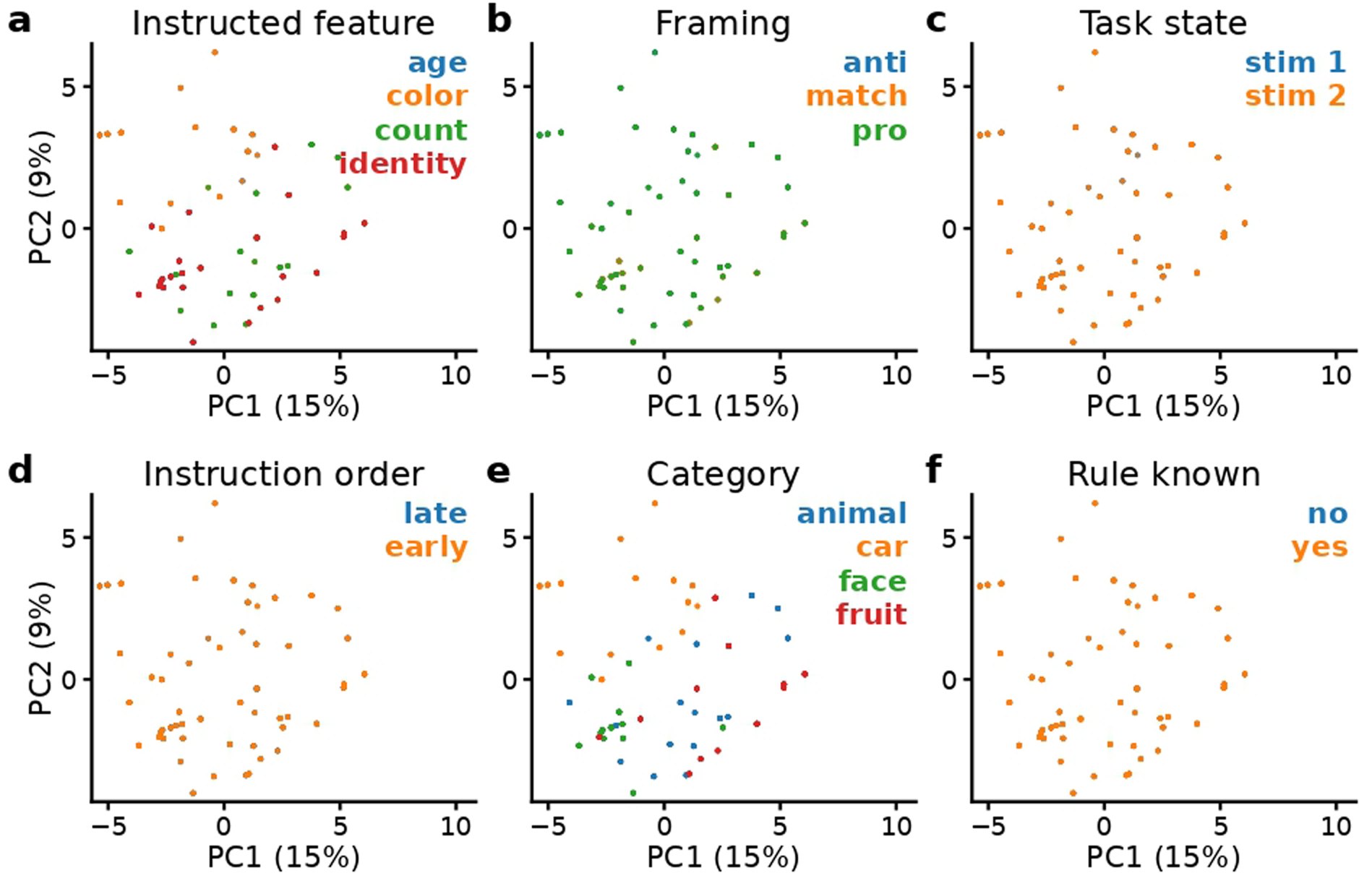
Representations from CLIP image embeddings, subsampled to 25% of trials for visualization. **a**, Two first principal components from CLIP embeddings across trials, separated by instructed feature. Variance explained between parentheses. **b**, Same, for task framing; CLIP embeddings unchanged. **c**, Same, for current task state (first vs. second stimulus; CLIP embeddings unchanged). **d**, Same, for instruction order; CLIP embeddings unchanged. **e**, Same, for stimulus category. **f**, Same, for whether the trial instruction was already known; CLIP embeddings unchanged.

**Extended Data Fig 7.**
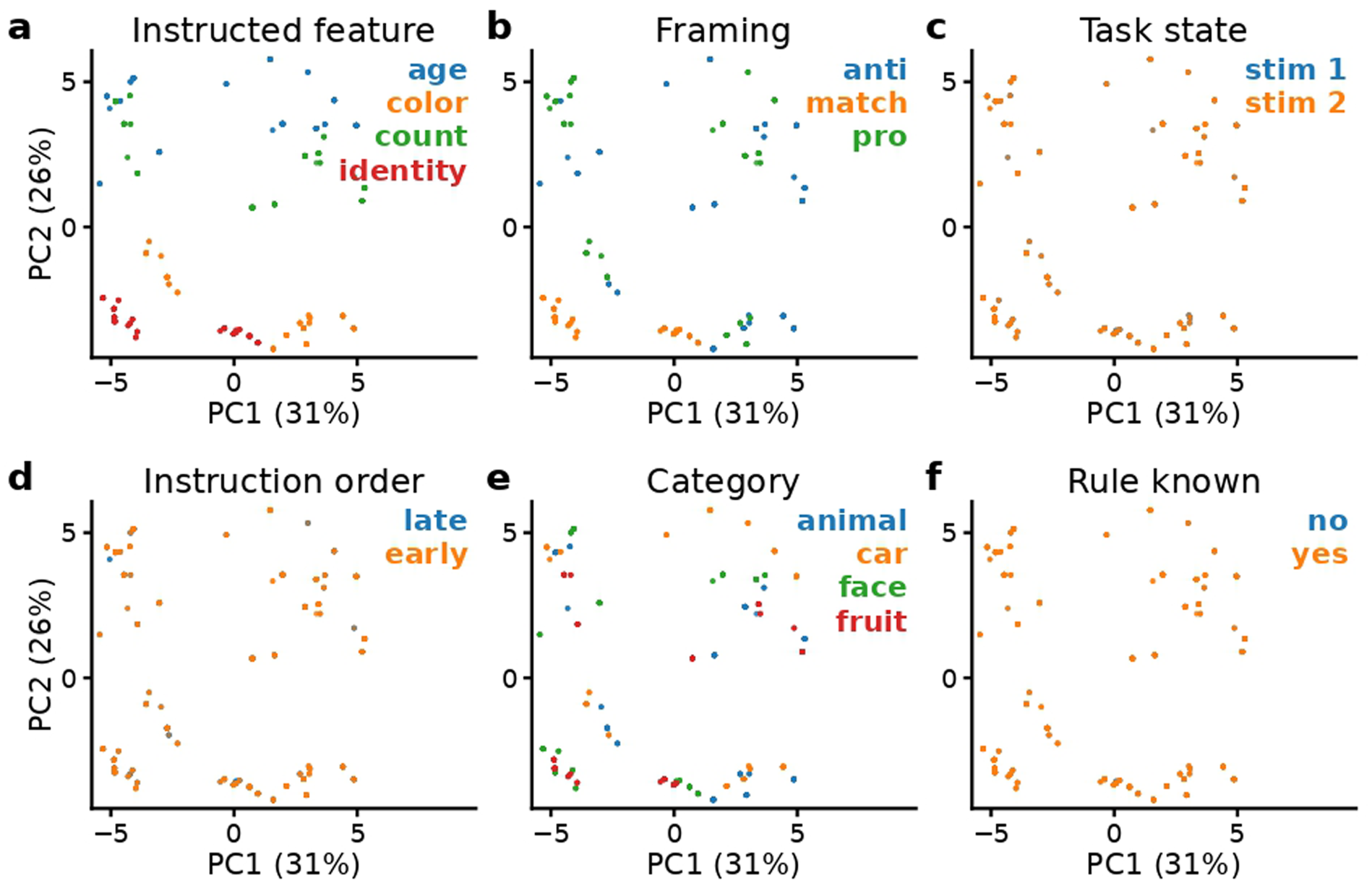
Representations from sBERT text embeddings, subsampled to 25% of trials for visualization. **a**, Two first principal components, from sBERT embeddings across trials, separated by instructed feature. Variance explained between parentheses. **b**, Same, for task framing. **c**, Same, for current task state (first vs. second stimulus; sBERT embeddings unchanged). **d**, Same, for instruction order; sBERT embeddings unchanged. **e**, Same, for stimulus category. **f**, Same, for whether the trial instruction was already known; sBERT embeddings unchanged.

**Extended Data Fig 8.**
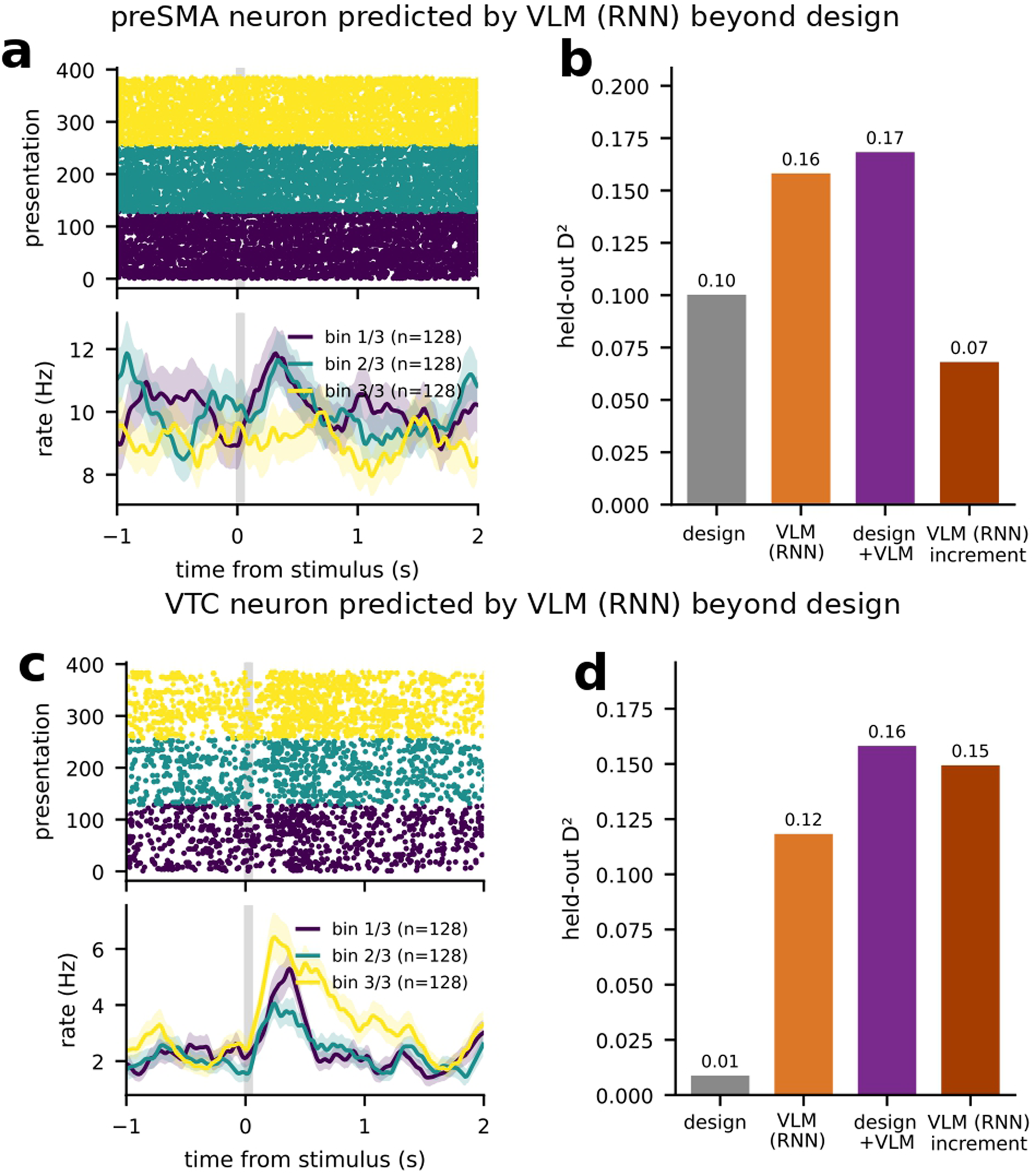
Examples of neurons which are predicted by the VLM’s task solution space beyond the experimental design factors. **a**, Raster plot and PSTH (shaded area indicates s.e.m.) for a preSMA neuron whose activity is explained by the VLM, plotted with respect to tertiles of VLM principal component 13. **b**, Poisson-Gamma’s ability to predict this neuron’s activity from the experimental design factors versus factors plus VLM, as well as the VLM’s increment. **c**, Same as a., for a VTC neuron plotted with respect to VLM principal component 6. **d**, Same as b., for the VTC neuron.

**Extended Data Fig 9.**
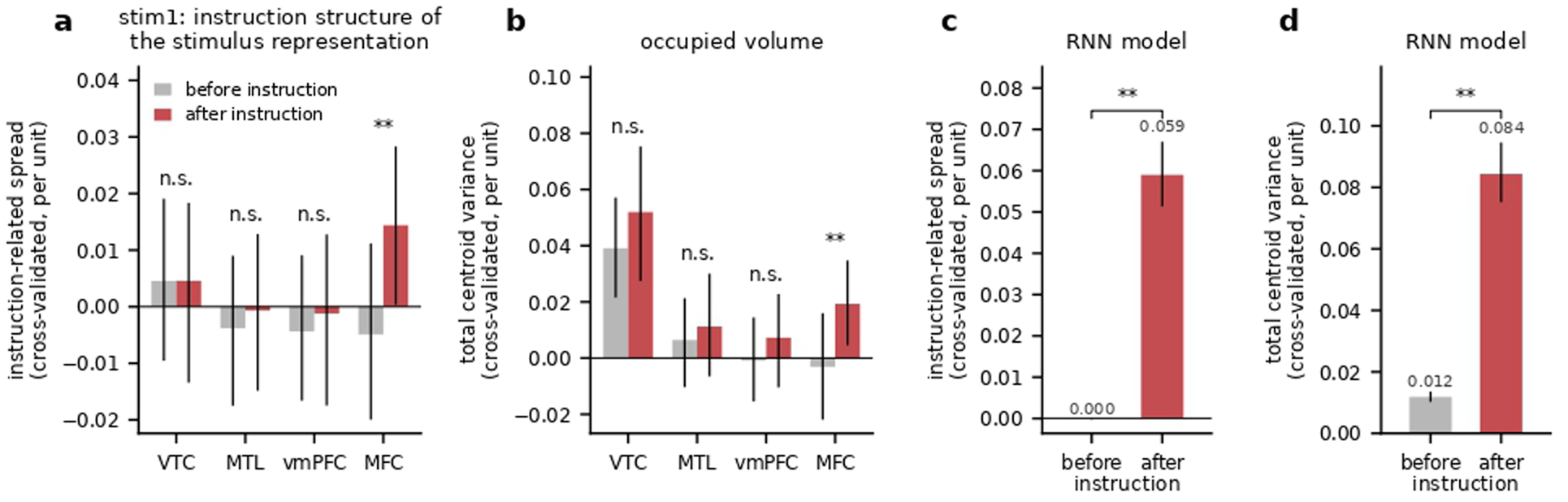
Instructions expand representation volume in RNNs and MFC. **a**, Comparing instruction-related spread of representations in neural space, during first stimulus presentation, before and after instruction presentations. **b**, Expansion of volume occupied by neural representations comparing pre- and post-instructions. **c**, Same as a., for the RNN module of the VLM. **d**, Same as b., for the RNN module of the VLM. Error bars indicate s.e.m.

**Extended Data Table 1.**
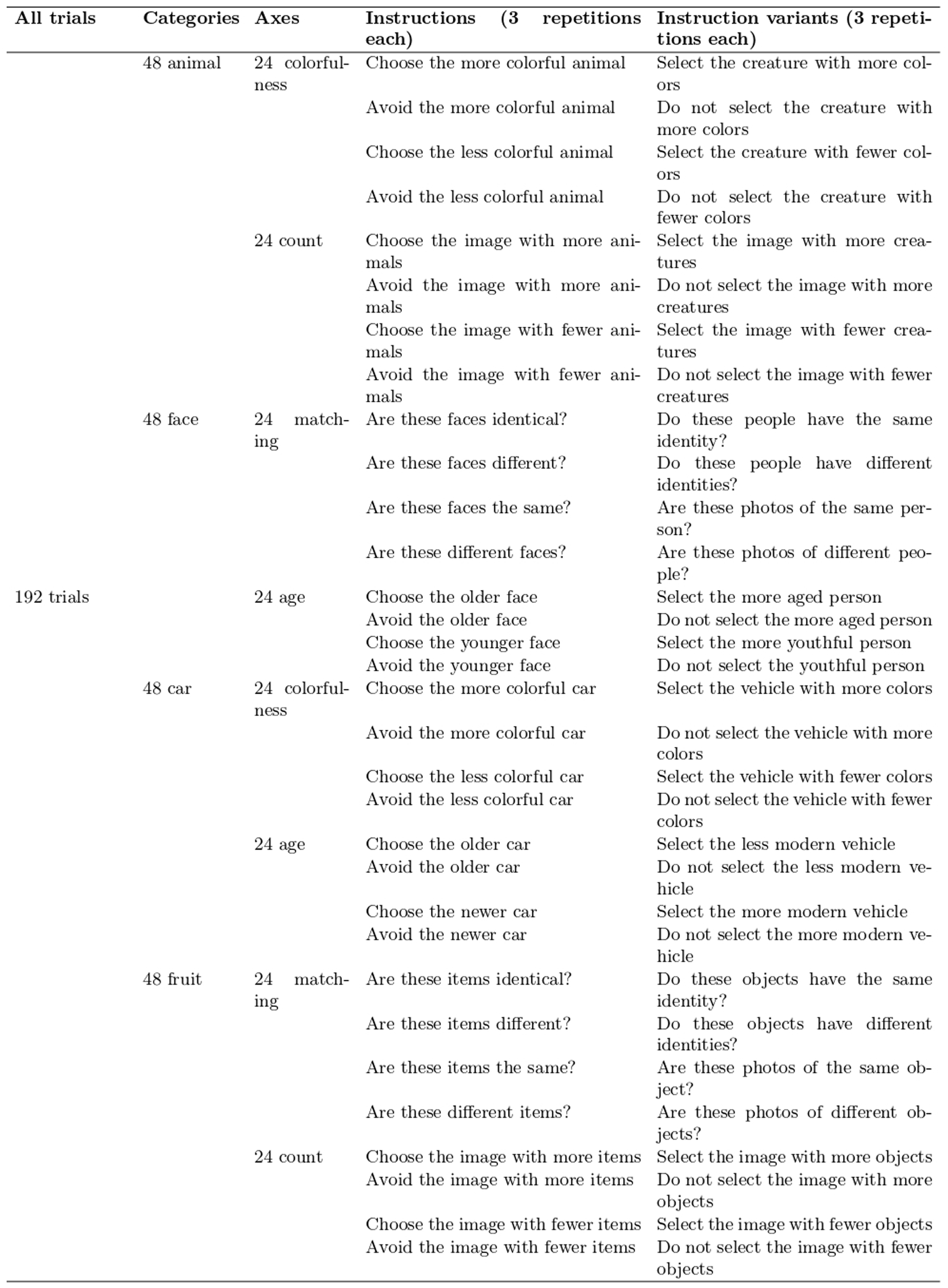
Verbal task instructions. Each of the 64 instructions was presented three times per session (192 trials).

| **All trials** | **Categories** | **Axes** | **Instructions (3 repetitions each)** | **Instruction variants (3 repeti-tions each)** |
|---|---|---|---|---|
| 192 trials | 48 animal | 24 colorful-ness | Choose the more colorful animal | Select the creature with more col-ors |
|  |  |  | Avoid the more colorful animal | Do not select the creature with more colors |
|  |  |  | Choose the less colorful animal | Select the creature with fewer col-ors |
|  |  |  | Avoid the less colorful animal | Do not select the creature with fewer colors |
|  |  | 24 count | Choose the image with more ani-mals | Select the image with more crea-tures |
|  |  |  | Avoid the image with more ani-mals | Do not select the image with more creatures |
|  |  |  | Choose the image with fewer ani-mals | Select the image with fewer crea-tures |
|  |  |  | Avoid the image with fewer ani-mals | Do not select the image with fewer creatures |
|  | 48 face | 24 match-ing | Are these faces identical? | Do these people have the same identity? |
|  |  |  | Are these faces different? | Do these people have different identities? |
|  |  |  | Are these faces the same? | Are these photos of the same per-son? |
|  |  |  | Are these different faces? | Are these photos of different peo-ple? |
|  |  | 24 age | Choose the older face | Select the more aged person |
|  |  |  | Avoid the older face | Do not select the more aged person |
|  |  |  | Choose the younger face | Select the more youthful person |
|  |  |  | Avoid the younger face | Do not select the youthful person |
|  | 48 car | 24 colorful-ness | Choose the more colorful car | Select the vehicle with more colors |
|  |  |  | Avoid the more colorful car | Do not select the vehicle with more colors |
|  |  |  | Choose the less colorful car | Select the vehicle with fewer colors |
|  |  |  | Avoid the less colorful car | Do not select the vehicle with fewer colors |
|  |  | 24 age | Choose the older car | Select the less modern vehicle |
|  |  |  | Avoid the older car | Do not select the less modern ve-hicle |
|  |  |  | Choose the newer car | Select the more modern vehicle |
|  |  |  | Avoid the newer car | Do not select the more modern ve-hicle |
|  | 48 fruit | 24 match-ing | Are these items identical? | Do these objects have the same identity? |
|  |  |  | Are these items different? | Do these objects have different identities? |
|  |  |  | Are these items the same? | Are these photos of the same ob-ject? |
|  |  |  | Are these different items? | Are these photos of different ob-jects? |
|  |  | 24 count | Choose the image with more items | Select the image with more objects |
|  |  |  | Avoid the image with more items | Do not select the image with more objects |
|  |  |  | Choose the image with fewer items | Select the image with fewer objects |
|  |  |  | Avoid the image with fewer items | Do not select the image with fewer objects |

**Extended Data Table 2.** Summary of patient demographics and single-neuron recordings across brain regions. Each entry indicates the total number of well-isolated neurons obtained from spike sorting. Values in parentheses correspond to left and right hemisphere counts, respectively.

| Session | Gender | Age | Seizure onset zone | VTC | HPC | AMY | preSMA | dACC | vmPFC |
|---|---|---|---|---|---|---|---|---|---|
| P81cs | F | 28 | Left mesial temporal | 25 (7, 18) | 9 (1, 8) | 38 (18, 20) | 33 (16, 17) | 9 (9, 0) | 22 (6, 16) |
| P82cs | M | 42 | Bilateral mesial temporal | 0 | 15 (2, 13) | 18 (14, 4) | 5 (5, 0) | 9 (7, 2) | 1 (1, 0) |
| P86cs | F | 39 | Not localized | 37 (23, 14) | 1 (0, 1) | 19 (3, 16) | 25 (20, 5) | 18 (7, 11) | 16 (5, 11) |
| P90cs | M | 32 | Left mesial temporal | 1 (0, 1) | 17 (14, 3) | 6 (5, 1) | 3 (3, 0) | 13 (5, 8) | 11 (11, 0) |
| P92cs | F | 30 | Left mesial occipital | 17 (8, 9) | 17 (4, 13) | 12 (6, 6) | 22 (8, 14) | 3 (3, 0) | 13 (0, 13) |
| P97cs | M | 32 | Right mesial temporal | 3 (0, 3) | 12 (7, 5) | 10 (5, 5) | 4 (0, 4) | 1 (1, 0) | 7 (1, 6) |
| P98cs | F | 30 | Bilateral mesial temporal | 21 (15, 6) | 22 (22, 0) | 4 (4, 0) | 8 (4, 4) | 0 | 18 (16, 2) |
| P99cs | F | 39 | Left lateral temporal | 2 (0, 2) | 13 (2, 11) | 1 (1, 0) | 17 (13, 4) | 6 (6, 0) | 7 (4, 3) |
| P100cs | M | 25 | Right mesial temporal | 18 (3, 15) | 6 (0, 6) | 12 (4, 8) | 18 (15, 3) | 0 | 12 (10, 2) |
| P102cs | F | 55 | Right mesial temporal | 32 (16, 16) | 15 (10, 5) | 12 (7, 5) | 20 (4, 16) | 15 (4, 11) | 11 (11, 0) |
| P103cs | M | 51 | Bilateral mesial temporal | 8 (6, 2) | 10 (2, 8) | 11 (7, 4) | 30 (18, 12) | 0 | 13 (9, 4) |

